# *Ehrlichia* Wnt short linear motif ligand mimetic deactivates the Hippo pathway to engage the anti-apoptotic Yap-GLUT1-BCL-xL axis

**DOI:** 10.1101/2023.03.06.531456

**Authors:** Caitlan D. Byerly, LaNisha L. Patterson, Nicholas A. Pittner, Regina N. Solomon, Jignesh G. Patel, Madison R. Rogan, Jere W. McBride

## Abstract

*Ehrlichia chaffeensis* TRP120 effector has evolved short linear motif (SLiM) ligand mimicry to repurpose multiple evolutionarily conserved cellular signaling pathways including Wnt, Notch and Hedgehog. In this investigation, we demonstrate that *E. chaffeensis* and recombinant TRP120 deactivate Hippo signaling resulting in activation of Hippo transcription coactivator Yap and target gene expression. Moreover, a homologous 6 amino acid (QDVASH) SLiM shared by TRP120 and Wnt3a/5a ligands phenocopied Yap and β-catenin activation induced by *E. chaffeensis,* rTRP120 and Wnt5a. Similar Hippo gene expression profiles were also stimulated by *E. chaffeensis,* rTRP120, SLiM and Wnt5a. Single siRNA knockdown of Hippo transcription co-activator/factors (Yap and TEAD) significantly decreased *E. chaffeensis* infection. Yap activation was abolished in THP-1 Wnt Frizzled-5 (Fzd5) receptor knockout cells (KO), demonstrating Fzd5 receptor dependence. In addition, TRP120 Wnt-SLiM antibody blocked Hippo deactivation (Yap activation). Expression of anti-apoptotic Hippo target gene *SLC2A1* (encodes glucose transporter 1; GLUT1) was upregulated by *E. chaffeensis* and corresponded to increased levels of GLUT1. Conversely, siRNA knockdown of *SLC2A1* significantly inhibited infection. Higher GLUT1 levels correlated with increased BCL-xL and decreased Bax levels. Moreover, blocking Yap activation with the inhibitor Verteporfin induced apoptosis that corresponded to significant reductions in levels of GLUT1 and BCL-xL, and activation of Bax and Caspase-3 and -9. This study identifies a novel shared Wnt/Hippo SLiM ligand mimetic and demonstrates that *E. chaffeensis* deactivates the Hippo pathway to engage the anti-apoptotic Yap-GLUT1-BCL-xL axis.

---

*Ehrlichia chaffeensis* is a Gram-negative, obligatory intracellular bacterium and the etiologic agent of the most prevalent and life-threatening tick-borne disease, human monocytic ehrlichiosis (HME). *Ehrlichia chaffeensis* preferentially infects mononuclear phagocytes, where it replicates within cytosolic, membrane-bound vacuoles and escapes host defenses through mechanisms executed by tandem repeat protein (TRP) effectors secreted by the type 1 secretion system (T1SS) (1). In the past decade, *E. chaffeensis* 120kDa tandem repeat protein (TRP120) has emerged as a model moonlighting effector that functions as a nucleomodulin, ubiquitin ligase and ligand mimetic to reprogram the mononuclear phagocyte and escape host innate immune defenses (2–4). TRP120 utilizes ligand mimicry to interact with various receptors to reprogram host cell signaling pathways conserved amongst eukaryotes, including Wnt, Notch and Hedgehog via novel tandem repeat <u>s</u>hort <u>li</u>near <u>m</u>otifs (SLiMs) within the TRP domain (5–9).

We previously demonstrated that *E. chaffeensis* activates canonical Wnt signaling by direct interaction of a TRP120 Wnt SLiM ligand mimetic and the host cell Wnt Frizzled 5 (Fzd5) receptor (6). Interaction between canonical Wnt ligands and Fzd5 receptor is known to stimulate Wnt transcriptional factor β-catenin but can also result in deactivation of Hippo signaling which coincides with the activation of transcription regulator Yes-associated protein (Yap) through Wnt-Hippo Fzd receptor crosstalk (10). SLiMs are short (3-11 amino acids) linear sequences typically found within intrinsically disordered protein domains that are responsible for mediating various cellular mechanisms through SLiM-protein interactions (11, 12). Interestingly, there are 23 predicted SLiMs in 14 proteins involved in Wnt signal transduction, including Axin, Dvl, and β-catenin (13). However, Wnt ligand SLiMs that mimic endogenous ligands, leading to pathway activation, have only recently been identified in *Ehrlichia* (6).

The Hippo pathway, discovered in *Drosophila* in 2003, is evolutionarily conserved in metazoans and universally recognized as a key regulator in embryogenesis, organ size, tissue homeostasis, cell proliferation, apoptosis, and tumorigenesis (14–18). Typically, when the Hippo pathway is active, the downstream transcriptional co-activator Yap is phosphorylated and deactivated, preventing nuclear translocation and activation of Hippo gene targets. When the Hippo pathway is activated, phosphorylation and deactivation of Yap occurs which in turn induces β-catenin deactivation and apoptosis (18, 19). Hippo pathway deactivation occurs when Wnt5a or Wnt3a ligands bind to the Wnt Fzd5 receptor, resulting in Yap and β-catenin activation and engagement of Hippo transcription factors TEAD and TCF, respectively (10, 20–24).

Regulation of apoptosis as a survival strategy is well-documented during *E. chaffeensis* infection (25). Mitochondria are the primary regulators of apoptosis by both intrinsic and extrinsic pathways, thus inhibition of mitochondrial outer membrane permeabilization (MOMP) is required to prevent apoptosis. It is known that *E. chaffeensis* stabilizes the mitochondria with effector Etf-1 by regulating mitochondrial matrix protein manganese superoxide dismutase (MnSOD) to induce antioxidative protection, thereby inhibiting apoptosis (25). Further, *E. chaffeensis* utilizes a TRP120 Hedgehog SLiM to activate Hedgehog signaling which prevents intrinsic apoptosis by maintaining BCL-2 levels and mitochondrial stability (9). However, there are likely other mechanisms *E. chaffeensis* engages to stabilize mitochondria such as modulation of other anti-apoptotic BCL-2 family of proteins, including BCL-xL, which is regulated by the Hippo pathway (26).

The Hippo pathway regulates various innate and metabolic responses including glycolysis and apoptosis (27–29). When Hippo signaling is deactivated, activated Yap binds TEAD and induces *SLC2A1* [encodes glucose transporter 1 (GLUT1)] upregulation, thereby promoting glycolysis and cell growth and apoptosis inhibition (26, 29–31). GLUT1 is a highly conserved glucose transporter that regulates glucose metabolism and prevents apoptosis by regulating the BCL-2 family of proteins. Specifically, GLUT1 amplifies anti-apoptotic BCL-xL levels and inhibits Bax and subsequent activation of caspases, resulting in an anti-apoptotic environment (26, 29, 32–35). In contrast, GLUT1 deficiency induces expression of pro-apoptotic proteins Bax, Bak, Bim and Bid, and inhibits expression of anti-apoptotic proteins MCL-1 and BCL-xL (36).

The Hippo pathway is well known for its role in cancer, but has recently been implicated in viral infections, including Hepatitis B virus (HBV), Hepatitis C virus (HCV), Human papillomavirus (HPV), Epstein-Barr virus (EBV), and Kaposi Sarcoma-associated herpesvirus (KSHV) (37). However, there are only a few reports of Hippo exploitation by parasites, fungi and bacteria (38–40). Yap activation by viruses has been reported, but the precise mechanism whereby deactivation of Hippo signaling occurs to activate Yap remains unclear (37). Studies have shown that Yap activation during HBV infection triggers hepatocarcinogenesis and pathogenesis of the liver and may cause HBV-induced hepatocellular carcinoma. Additionally, HBV infection of Alb-preΔS2 transgenic mice increases expression of Hippo target genes *BIRC5, ANKRD1, CTGF*, and *CYR61* (37, 41). HPV E6 major oncoprotein inhibits active Yap degradation, and Yap knockdown impairs E6-mediated cell proliferation indicating that Yap activation plays a role in the proliferation of cervical cancer cells (42).

Although the Hippo pathway is targeted by multiple pathogens, the pathogen-host interactions and mechanisms involved in Hippo pathway exploitation have not been defined. We have previously identified an *Ehrichia* SLiM that activates Wnt signaling. Therefore, since Hippo signaling is initiated through Wnt Fzd receptors we considered that Hippo signaling may be regulated through the same ligand-receptor complex during infection (43). This investigation reveals a strategy whereby *E. chaffeensis* utilizes a eukaryotic Wnt SLiM ligand motif interaction with the Fzd5 receptor to deactivate Hippo signaling, thereby activating the Yap-GLUT1-BCL-xL axis to promote an anti-apoptotic cellular environment.

## Results

### *E. chaffeensis* activates Yap and Hippo gene expression

Hippo deactivation mediated by Wnt ligand engagement of the Fzd5 receptor results in Yap activation and nuclear translocation, where it binds the transcription factor TEAD to regulate Hippo gene targets (10, 18, 20, 24, 44). Recent studies demonstrate that *E. chaffeensis* directly interacts with Fzd5 receptor to activate β-catenin. To investigate whether *E. chaffeensis* activates Yap via Hippo-Wnt ligand-receptor crosstalk, we detected active Yap in the nucleus of infected THP-1 cells within 4 h post-infection (hpi). Moreover, progressive nuclear accumulation of active Yap was observed over 48 hpi compared to uninfected controls (**Fig. 1A, C, D**). Further, active Yap accumulated in the nucleus in *E. chaffeensis*-infected primary human monocytes (10 hpi) compared to the uninfected control, providing further evidence of *E. chaffeensis*-mediated Hippo deactivation (**Fig. 1B, E**). To further examine the role of *E. chaffeensis* in Hippo deactivation (Yap activation), we examined Hippo pathway gene transcription using a human Hippo signaling PCR array **(Fig. 1F)**. Significant activation of Hippo pathway component genes was detected during *E. chaffeensis* infection, with the majority (63%) of Hippo genes being upregulated, including major Hippo and Wnt components *YAP*, *TAZ*, *TEAD1*, *TEAD2, TEAD3, TEAD4,* and *DVL2* compared to controls **(Fig. 1F)**.

**Fig. 1.**
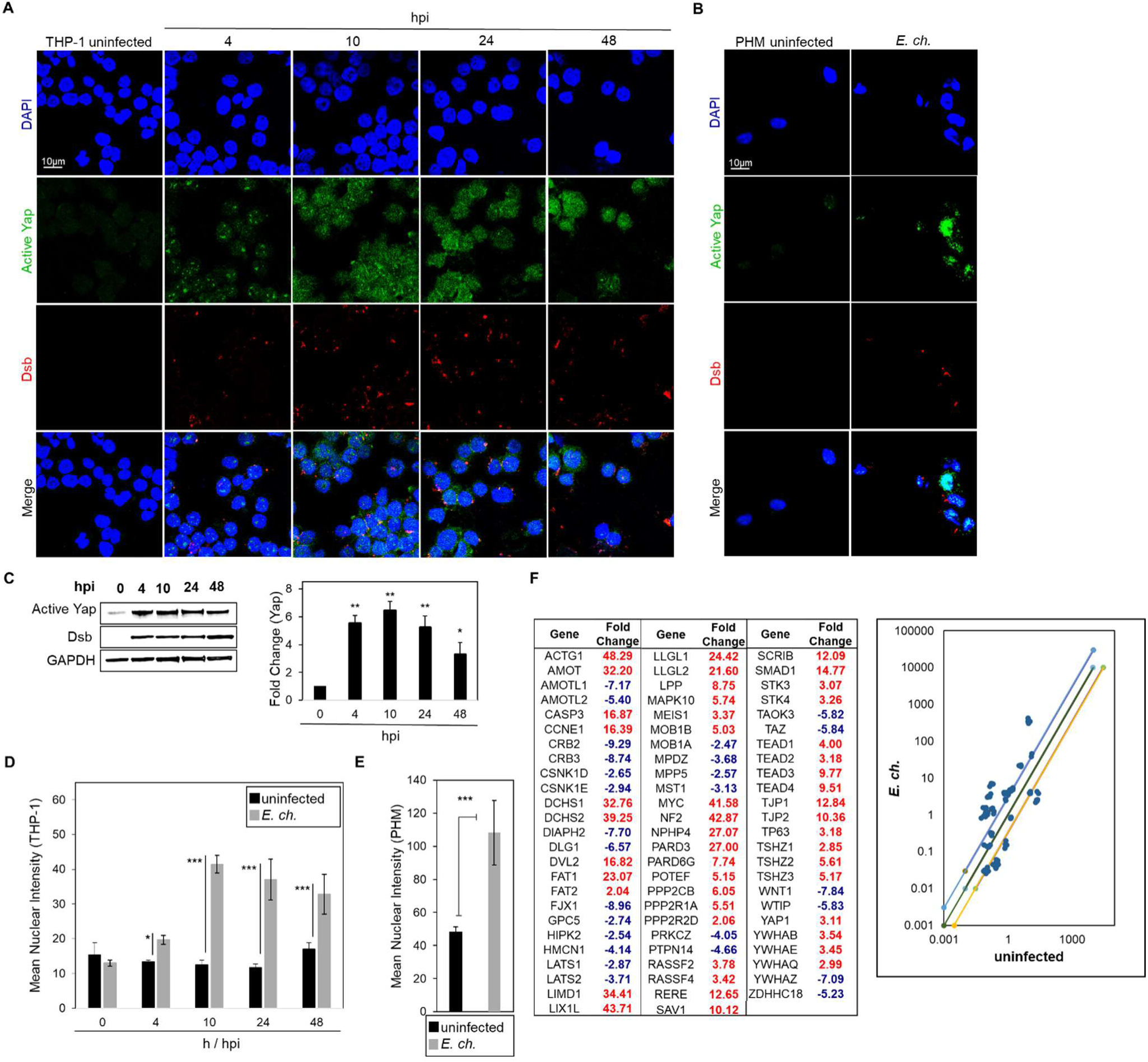
*E. chaffeensis* activates Yap and Hippo gene expression. (A) Confocal immunofluorescence micrographs showing temporal Yap activation (green) at 0, 4, 10, 24 and 48 h post-infection (hpi) in *E. chaffeensis*-infected THP-1 cells. Anti-Dsb antibody (red) confirms *E. chaffeensis* infection (scale bar = 10 μm). (B) Confocal immunofluorescence micrographs showing Yap activation (green) in uninfected and *E. chaffeensis-*infected (10 h) primary human monocytes. Anti-Dsb antibody (red) confirms *E. chaffeensis* infection (scale bar = 10 μm). (A-B) Experiments were performed with three biological and technical replicates. Randomized areas/slide (n=10) were selected to detect active Yap. (C) Western blot analysis depicting active Yap levels at 0, 4, 10, 24 and 48 hpi with GAPDH as a loading control. Anti-Dsb antibody demonstrates *E. chaffeensis* infection. Bar graph (right) represents densitometry values of Western blot normalized to GAPDH. Western blots were performed with three biological and technical replicates for *t*-test analysis. Data are represented as means ± SD (\**p*< 0.05; \*\**p*< 0.01). (D-E) Intensity graphs demonstrate the mean nuclear accumulation of active Yap in THP-1 cells and primary human monocytes, respectively. Analysis was performed using ImageJ to determine mean grey value from randomized areas/slide (n=10) and data shown as mean ± SD (\**p*< 0.05; \*\*\**p*< 0.001). (F) Table represents normalized expression of significantly regulated Hippo array genes between *E. chaffeensis-*infected and uninfected cells at 24 h. The scatterplot represents the expression of all Hippo array genes. The top and bottom lines depict a 2-fold upregulation or downregulation, respectively, compared to uninfected control. Scatterplots are representative of three (*n*=3) biological and technical replicates.

### TRP120 activates Yap and Hippo gene expression

To further examine the role of TRP120 in Hippo deactivation, THP-1 cells and primary human monocytes were incubated with recombinant TRP120 protein (rTRP120-FL), and Yap activation was examined using confocal microscopy **(Fig. 2A-B)**. Active Yap accumulated in the nucleus of THP-1 cells **(Fig. 2A, C)** and primary human monocytes **(Fig. 2B, D)** at 6 and 10 h post-treatment (hpt), respectively, consistent with Yap activation by recombinant Wnt5a (rWnt5a). To further confirm the role of TRP120 in Hippo regulation, cells were stimulated with rTRP120-FL or rWnt5a for 24 h, and a transcriptional analysis was performed **(Fig. 2E-F)**. Hippo genes (45%) were significantly upregulated, including genes important for Hippo and Wnt signaling (*YAP, TAZ, TEAD4,* and *DVL2*) and 16% were downregulated **(Fig. 2E)**. In comparison, cells treated with rWnt5a had significant transcriptional upregulation of Hippo genes (65%) including *YAP*, *TAZ*, *TEAD1*, *TEAD2, TEAD3, TEAD4, WNT1* and *DVL2*, and 22% of genes were significantly downregulated (**Fig. 2F**). Though there were differential expression patterns of genes in TRP120 and Wnt5a treated cells, we found that 34 Hippo target genes including *YAP, TAZ, TEAD4,* and *DVL2* were upregulated in both rTRP120-FL and rWnt5a treatment. Together these data demonstrate that TRP120 independently and efficiently activates Yap to regulate Hippo target genes.

**Fig. 2.**
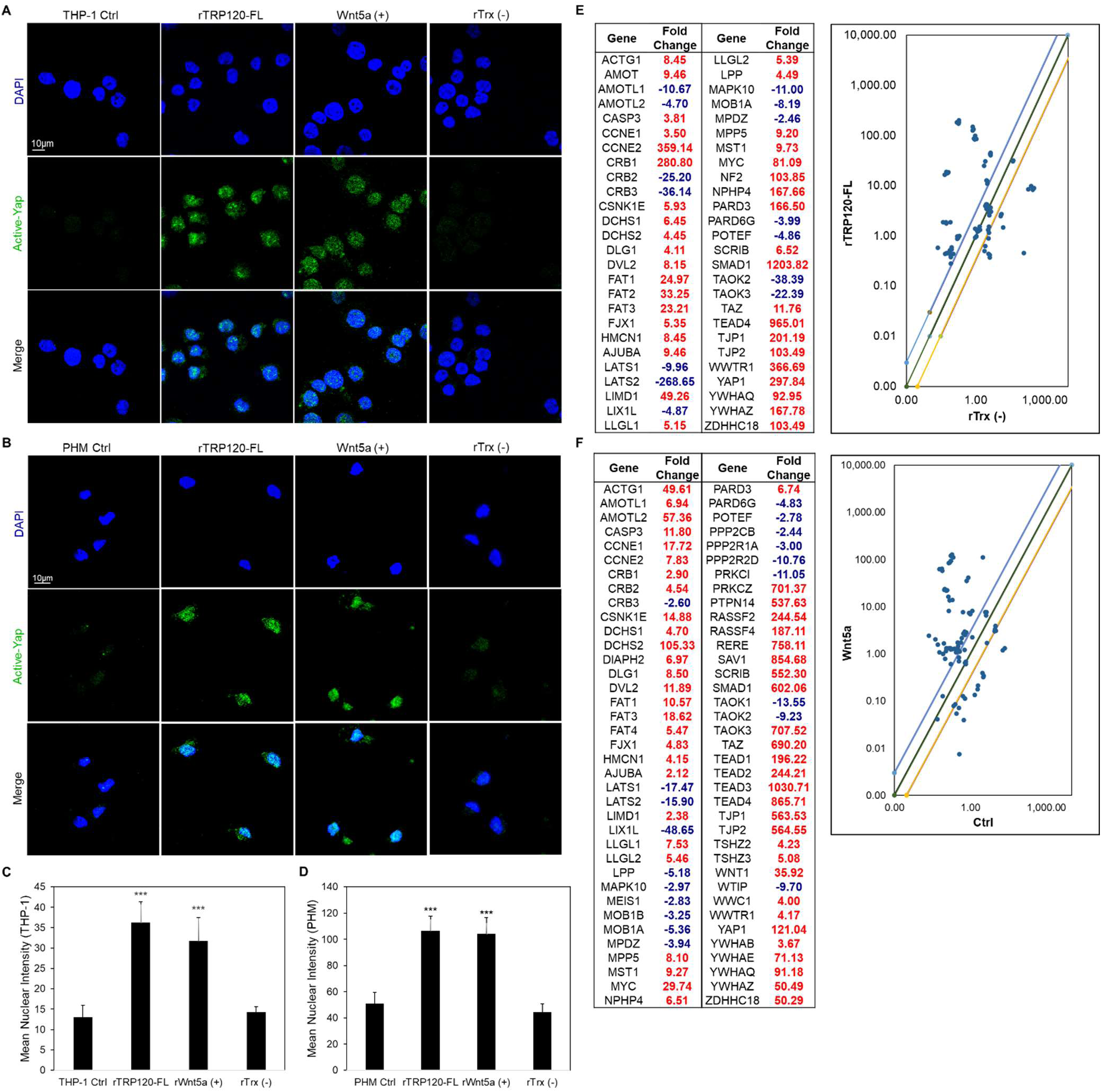
TRP120 activates Yap and Hippo gene targets. (A) Confocal immunofluorescence micrographs demonstrating rTRP120-FL-, rTrx-(−), rWnt5a-treated (+) (1 μg/mL) or untreated (control) THP-1 cells stained with active Yap antibody (green) 6 h post-treatment (hpt) (scale bar = 10 μm). (B) Confocal immunofluorescence microscopy of untreated (control) or rTRP120-FL-, rTrx-(−), rWnt5a-treated (+) (1 μg/mL) primary human monocytes stained with active Yap antibody (green) 10 hpt (scale bar = 10 μm). (A-B) Experiments were performed with three biological and technical replicates. Randomized areas/slide (n=10) were selected to detect active Yap. (C-D) Intensity graphs demonstrate the mean nuclear accumulation of active Yap in respective THP-1 cells and primary human monocytes. Analysis was performed using ImageJ to determine mean grey value from randomized areas/slide (n=10), and data shown as mean ± SD (\*\*\**p*< 0.001). (E) The table represents significantly regulated Hippo signaling PCR array gene expression in THP-1 cells stimulated with rTRP120-FL (1 μg/mL) after normalization to control cells treated with rTrx (1 μg/mL) at 24 h. The respective normalized expression of rTRP120-FL regulated Hippo array genes was performed with three biological and technical replicates. (F) The table represents significantly regulated Hippo signaling PCR array gene expression in THP-1 cells stimulated with rWnt5a (1 μg/mL) after normalization to DMSO-treated cells (control). The respective normalized expression of rWnt5a regulated Hippo array genes is representative of three biological replicates. (E-F) The scatterplot represents the expression of all Hippo array genes. The top and bottom scatterplot lines depict a 2-fold upregulation or downregulation, respectively, compared to control. Data is representative of three independent experiments (*n*=3).

### TRP120 Wnt SLiM regulates Hippo signaling

TRP120 contains a tandem repeat domain (TRD), with four tandem repeats, flanked by N- and C-terminal domains. Various TRP120 SLiMs have been reported within the TRD, and C-terminus that are relevant to *E. chaffeensis* pathobiology, including posttranslational modification motifs, DNA-binding motifs, and ubiquitin ligase catalytic motifs (6). We previously reported that *E. chaffeensis* TRP120 TRD utilizes SLiMs to regulate Wnt, Notch and Hedgehog signaling pathways (6, 8, 9). A TRP120 Wnt SLiM that activates Wnt signaling was previously reported and homology was identified between TRP120 and Wnt5a (6). However, based on a revised BLAST analysis of TRP120 with both Wnt5a and Wnt3a **(Fig. 3A)**, we identified a shorter Wnt SLiM (QDVASH) shared by both ligands (60% and 83% similarity, respectively) within the previously identified TRP120 Wnt-SLIM (IKDL<u>QDVASH</u>ESGVSDQ).

**Fig. 3.**
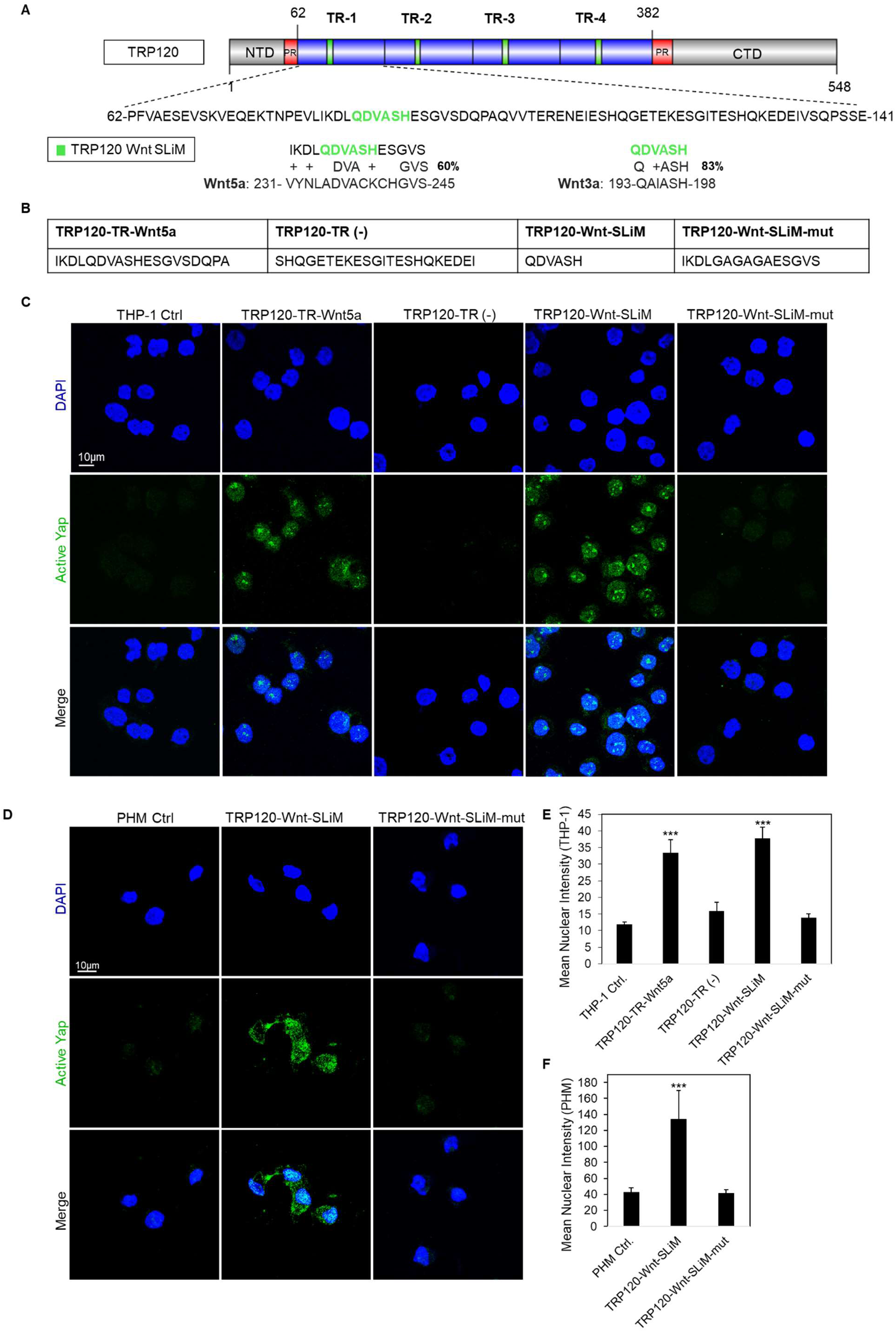
TRP120 Wnt SLiM regulates Hippo signaling. (A) Schematic representation of TRP120 showing domain organization; N-terminal (NTD), C-terminal (CTD), tandem repeat (TR1 – 4; 80 aa each) and partial repeat (PR) domain (70). A 6 amino acid short linear motif (SLiM) of high sequence similarity was identified using NCBI BLAST between the TRP120 TR and Wnt3a/Wnt5a ligands (activators of Yap) amino acid sequences. The complete amino acid sequence of one TR is shown with homologous Wnt SLiM identified in green and percent homology right of the sequence. (B) The table displays the various TRP120 peptide amino acid sequences used in the TRP120 Wnt SLiM study. TRP120-Wnt-SLiM represents the homology sequence identified though BLAST. TRP120-Wnt-SLiM-mut contains glycine and alanine substitutions in the Wnt SLiM region and is used as a negative control. TRP120-TR-Wnt5a is a 19 amino acid sequence that contains the identified TRP120 Wnt homology sequence. TRP120-TR (−) is a sequence within the TRP120-TR that does not contain the defined TRP120 Wnt homology sequence. (C) Confocal immunofluorescence microscopy of untreated (−) or peptide-treated THP-1 cells (1 μg/mL). THP-1 cells were stained with active Yap antibody and the micrograph shows increased levels of active Yap (green) in TRP120-TR-Wnt5a and TRP120-Wnt-SLiM-treated, but not in untreated, TRP120-TR (−) or TRP120-Wnt-SLiM-mut treated THP-1 cells (6 hpt)(scale bar = 10 μm). (D) Confocal immunofluorescence microscopy of untreated or SLiM/SLiM mutant peptide-treated primary human monocytes (10 h). The TRP120-Wnt-SLiM sequence upregulates active Yap (green) in primary human monocytes, but the corresponding mutant sequence does not (scale bar = 10 μm). (C-D) Experiments were performed with three biological and technical replicates. Randomized areas/slide (n=10) were selected to detect active Yap nuclear translocation. (E-F) Intensity graphs demonstrate the mean nuclear accumulation of active Yap in respective THP-1 cells and primary human monocytes. Analysis was performed using ImageJ to determine mean grey value from randomized areas/slide (n=10). Data are represented as means ± SD (\*\*\**p*< 0.001).

To investigate Wnt SLiM activation of Yap, THP-1 cells were treated for 6 h with two peptides that contained the following sequences: TRP120-Wnt-SLiM (6 aa) and TRP120-TR-Wnt5a (19 aa); and two control peptides that did not contain Wnt SLiM: TRP120-Wnt-SLiM-mut (15 aa; glycine/alanine substitutions) and TRP120-TR (−) (22 aa; TR sequence null of Wnt SLiM) **(Fig. 3B)** and Yap activation was determined **(Fig. 3C, E)**. Both TRP120-TR-Wnt5a and TRP120-Wnt-SLiM treatments stimulated significant Yap activation compared to TRP120-TR (−) and TRP120-Wnt-SLiM-mut controls **(Fig. 3E)**. Similarly, TRP120-Wnt-SLiM treatment significantly stimulated active Yap in primary human monocytes (10 hpt) compared to the TRP120-Wnt-SLiM-mut **(Fig. 3D, F)**. Additionally, to determine whether the newly defined TRP120-Wnt SLiM activates Wnt signaling, we treated THP-1 cells with TRP120-Wnt-SLiM and TRP120-Wnt-SLiM-mut and measured active β-catenin nuclear accumulation **(Fig. S1A)**. TRP120-Wnt-SLiM was able to significantly stimulate active β-catenin consistent with *E. chaffeensis* and Wnt5a, confirming that the 6 aa Wnt SLiM was completely responsible for Wnt signaling activation and the amino acids flanking the 6 aa Wnt SLiM are not significant.

Further, we treated THP-1 cells with a single amino acid TRP120-Wnt-SLiM histidine deletion mutant (QDVAS) and observed no significant activation of Yap or β-catenin, indicating that the 6 amino acid TRP120-Wnt-SLiM containing histidine is essential for activation **(Fig. S2)**. Various studies have demonstrated the importance of histidine in protein-protein interactions. In fact, histidine is known as the most active and versatile amino acid, is often the key residue in enzyme catalytic reactions, and is essential for protein interactions (45).

### TRP120 Wnt SLiM concentration dependent Hippo gene activation

To investigate if the TRP120-Wnt-SLiM regulates Hippo target genes, THP-1 cells were treated with TRP120-Wnt-SLiM (10 or 1000 ng/mL) and significant Hippo gene activation was detected in a concentration-dependent manner **(Fig. 4A-C)**. TRP120-Wnt-SLiM influenced Hippo gene expression, including Hippo and Wnt target genes *YAP*, *WNT1*, *DVL2*, *TEAD1*, and *TEAD4* consistent with *E. chaffeensis*, TRP120-, and WNT5a (10 ng/mL). Moreover, all Hippo genes were significantly upregulated in response to 100-fold higher TRP120-Wnt-SLiM concentration (1000 ng/mL). These data demonstrate that the defined TRP120-Wnt-SLiM activates Yap and regulates Hippo gene expression in a concentration dependent manner.

**Fig. 4.**
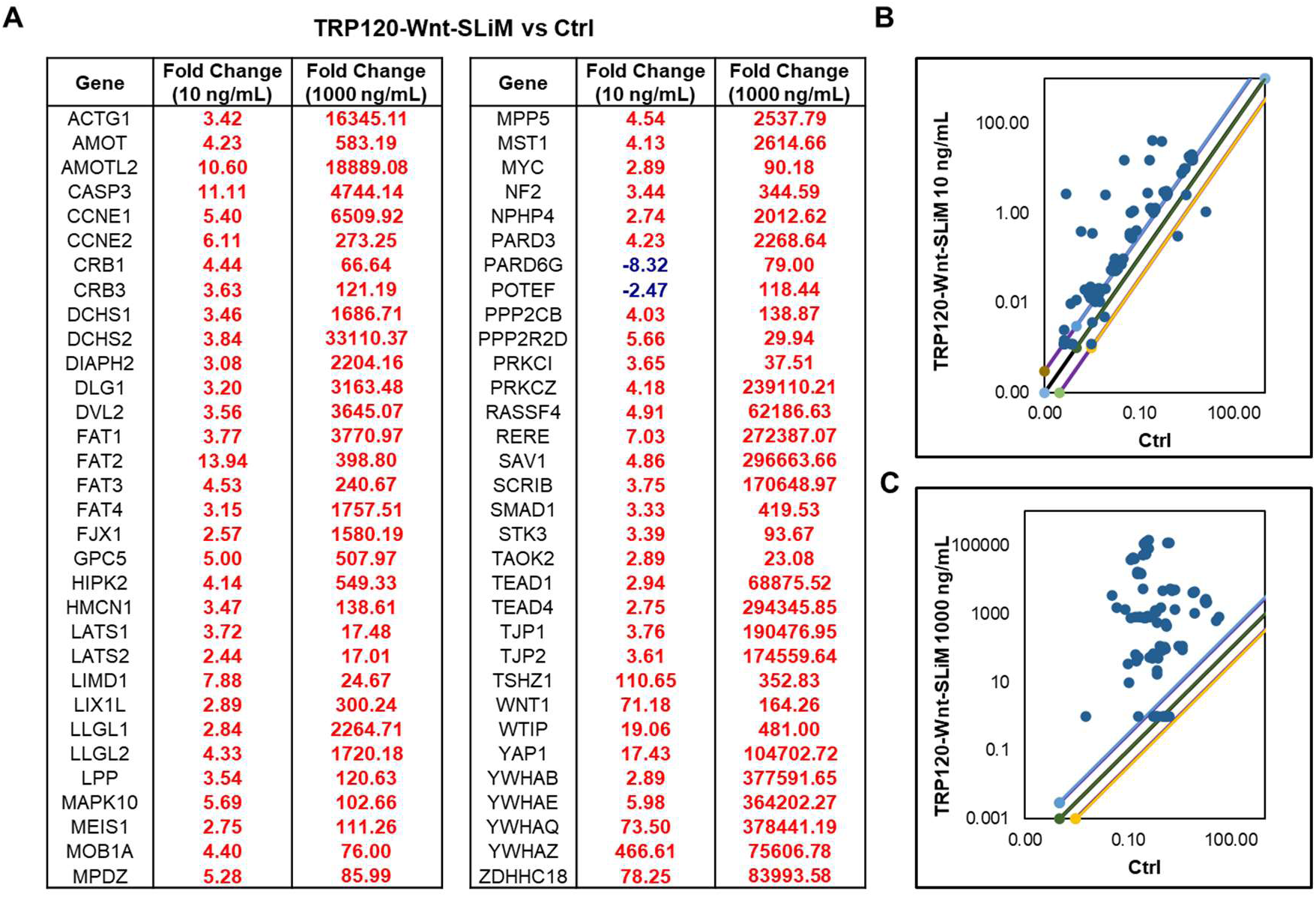
TRP120 Wnt SLiM concentration dependent Hippo gene activation. (A) Hippo signaling PCR array used for the analysis of the expression of 84 Hippo genes. THP-1 cells were treated with TRP120-Wnt-SLiM (10 and 1000 ng/mL) or left untreated (negative control) and were harvested at 24 h. The tables represent significant fold change in gene expression in TRP120-Wnt-SLiM-treated cells compared to untreated cells at respective concentrations. Data represents three (n=3) biological replicates. (B-C) The scatterplots represent the expression of all Hippo array genes. The top and bottom scatterplot lines depict a 2-fold upregulation or downregulation, respectively, compared to control. Scatterplots are representative of three independent experiments (*n*=3).

### Hippo co-activator and transcription factors influence infection

Although the Hippo pathway is widely recognized for its role in embryogenesis and tumorigenesis, it also plays a key role in regulating apoptosis, which is crucial for successful ehrlichial intracellular infection (3, 9, 14, 15, 25). To determine whether *E. chaffeensis* survival depends on Hippo transcriptional components, we used RNAi to individually silence genes for *YAP*, *TEAD1*, *TEAD3* and *TEAD4* (**Fig. 5A**). Ehrlichial load was significantly reduced in all transfection groups 24 h post RNAi transfection compared to the scramble siRNA-transfected controls **(Fig. 5B**).

**Fig. 5.**
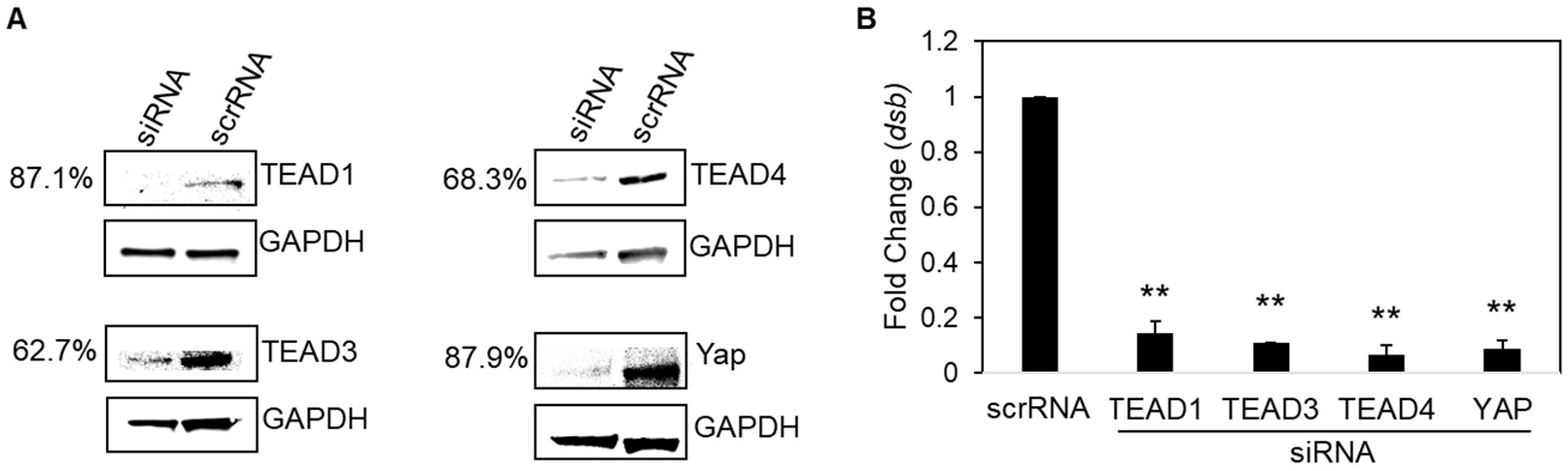
Hippo co-activator and transcription factors influence infection. (A) Western blots depict knockdown efficiency of small interfering RNA-transfected (siRNA) THP-1 cells, with scrambled siRNA (scrRNA) transfected THP-1 cells as control from whole-cell lysates (24 hpt). siRNA knockdown (%) indicates total percent knockdown of protein of interest relative to control, normalized to GAPDH. (B) THP-cells (24 hpt) were infected with *E. chaffeensis* (MOI 100) and harvested 24 hpi. Infected scrRNA cells are represented as positive control. qPCR amplification of the ehrlichial disulfide bond formation protein (*dsb*) gene was used to quantify *E. chaffeensis* infection. siRNA knockdown of Hippo transcription components TEAD1-4 and Yap significantly inhibits *E. chaffeensis* infection in THP-1 transfected cells. All knockdowns were performed with three biological and technical replicates for *t*-test analysis. Data are represented as mean ± SD (\*\**p*< 0.01).

### A TRP120-Wnt domain targeted antibody blocks Yap activation

To further elucidate the role of TRP120-Wnt-SLiM during *E. chaffeensis* infection, we investigated whether blocking *E. chaffeensis* infection or the TRP120-Wnt-SliM with a TRP120-Wnt-SliM-targeted antibody would inhibit Yap activation. *E. chaffeensis* infected and TRP120-Wnt-SliM treated cells in the presence of α-TRP120-Wnt-SliM demonstrated significant reduction in active Yap relative to *E. chaffeensis-*infected and TRP120-Wnt-SliM-treated cells in the presence of α-TRP120-PIS antibody (control) **(Fig. 6A-C)**. These data confirm that the TRP120-Wnt-SliM activates Yap and the SliM ligand activity can be blocked by antibody.

**Fig. 6.**
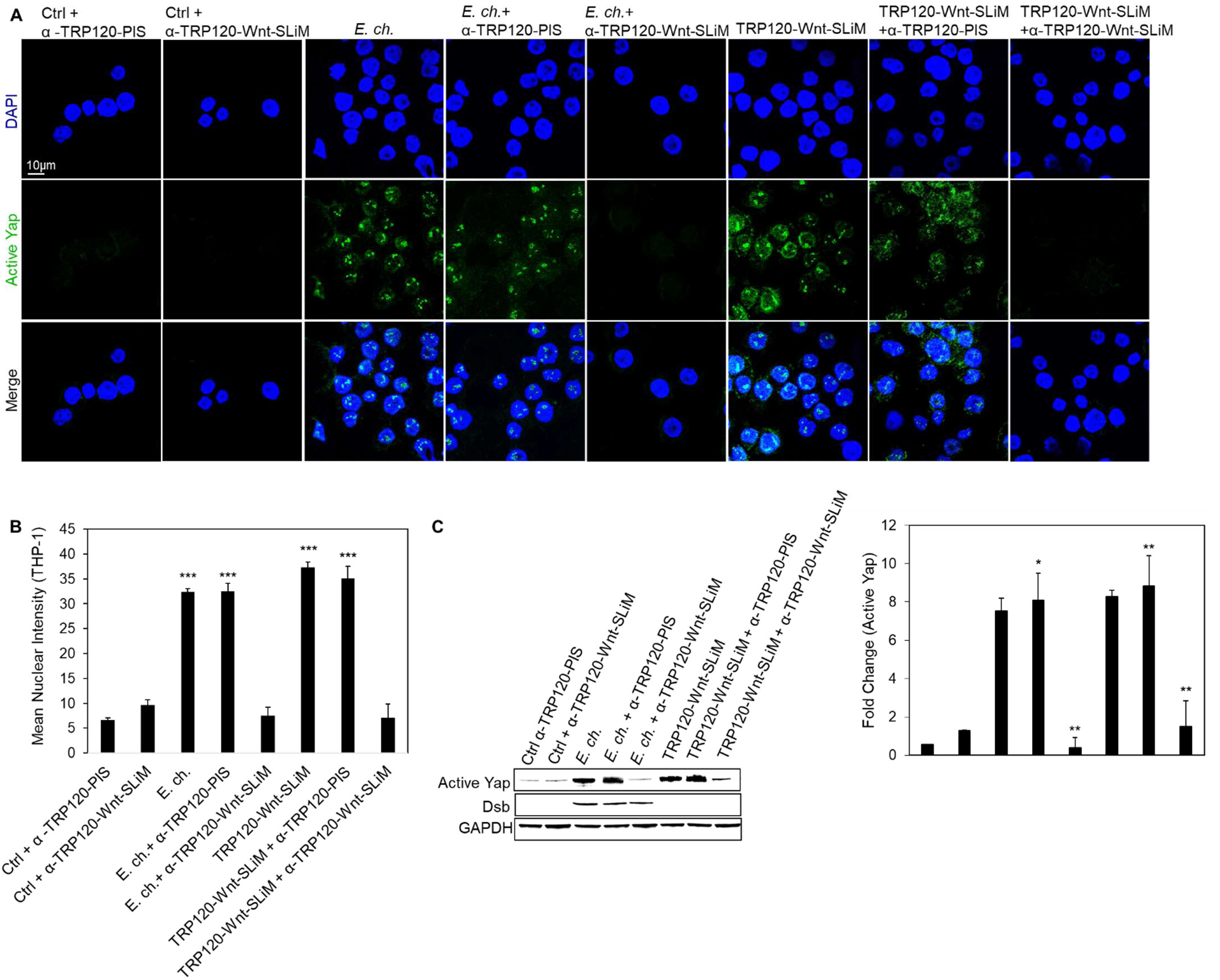
A TRP120-Wnt domain targeted antibody blocks Yap activation. (A) *E. chaffeensis* (MOI 100) and TRP120-Wnt-SLiM (1 μg/mL) were incubated with α-TRP120-Wnt-SLiM (targets TRP120 sequence DL<u>QDVASH</u>ESGVSDQPAQV) or α-TRP120-PIS (neg ctrl) (1.5 μg/mL) for 1 h or overnight, respectively, before incubation with THP-1 cells. THP-1 cells were harvested 6 hpt, immunostained with active Yap antibody (green), and visualized by confocal fluorescence microscopy (scale bar = 10 μm). Randomized areas/slide (n=10) were selected to detect active Yap nuclear translocation. (B) Intensity graph demonstrates the mean nuclear accumulation of active Yap in respective THP-1 cells. Analysis was performed using ImageJ mean grey value from randomized areas/slide (n=10). (C) Western blot analysis of treatment groups with GAPDH as a loading control with bar graph of Western blot analyzed from densitometry values normalized to GAPDH (A-C); α-TRP120-Wnt-SLiM inhibits active Yap upregulation in cells with *E. chaffeensis* or TRP120-Wnt-SLiM compared to α-TRP120-PIS. Untreated cells were incubated with α-TRP120-Wnt-SLiM or α-TRP120-PIS as negative controls. Experiments were performed with three biological and technical replicates and significance was determined through *t*-test analysis. Data are represented as means ± SD (\**p*< 0.05; \*\**p*< 0.01; \*\*\**p*< 0.001).

### Hippo deactivation is dependent on the Fzd5 receptor

During Hippo-Wnt receptor crosstalk, Wnt5a and Wnt3a ligands bind the Fzd5 receptor to deactivate Hippo signaling and activate Yap to engage Hippo gene transcription. We previously demonstrated that *E. chaffeensis* TRP120 TRD directly binds Fzd5 receptor (6). To determine the basis of this interaction regarding Hippo signaling, Fzd5 receptor knockout (KO) cells were used to determine the role of Fzd5 receptor in Yap activation. Fzd5 receptor KO or normal THP-1 cells (control) were infected with *E. chaffeensis* or treated with TRP120-Wnt-SliM. Fzd5 receptor KO cells exhibited no significant Yap activation compared to the control **(Fig. 7A-C)**. These results demonstrate that *E. chaffeensis* engages Fzd5 receptor to deactivate Hippo and activate Yap. Similarly, we determined that there was significant deactivation of β-catenin in THP-1 Fzd5 receptor KO cells infected with *E. chaffeensis* or treated with TRP120-Wnt-SliM, revealing that *E. chaffeensis* interacts with Fzd5 receptor to activate Wnt signaling **(Fig. S1B)**. However, unlike Yap, there was significant activation of β-catenin compared to control cells, likely due to contribution of other Fzd receptors known to interact with TRP120 (6).

**Fig 7.**
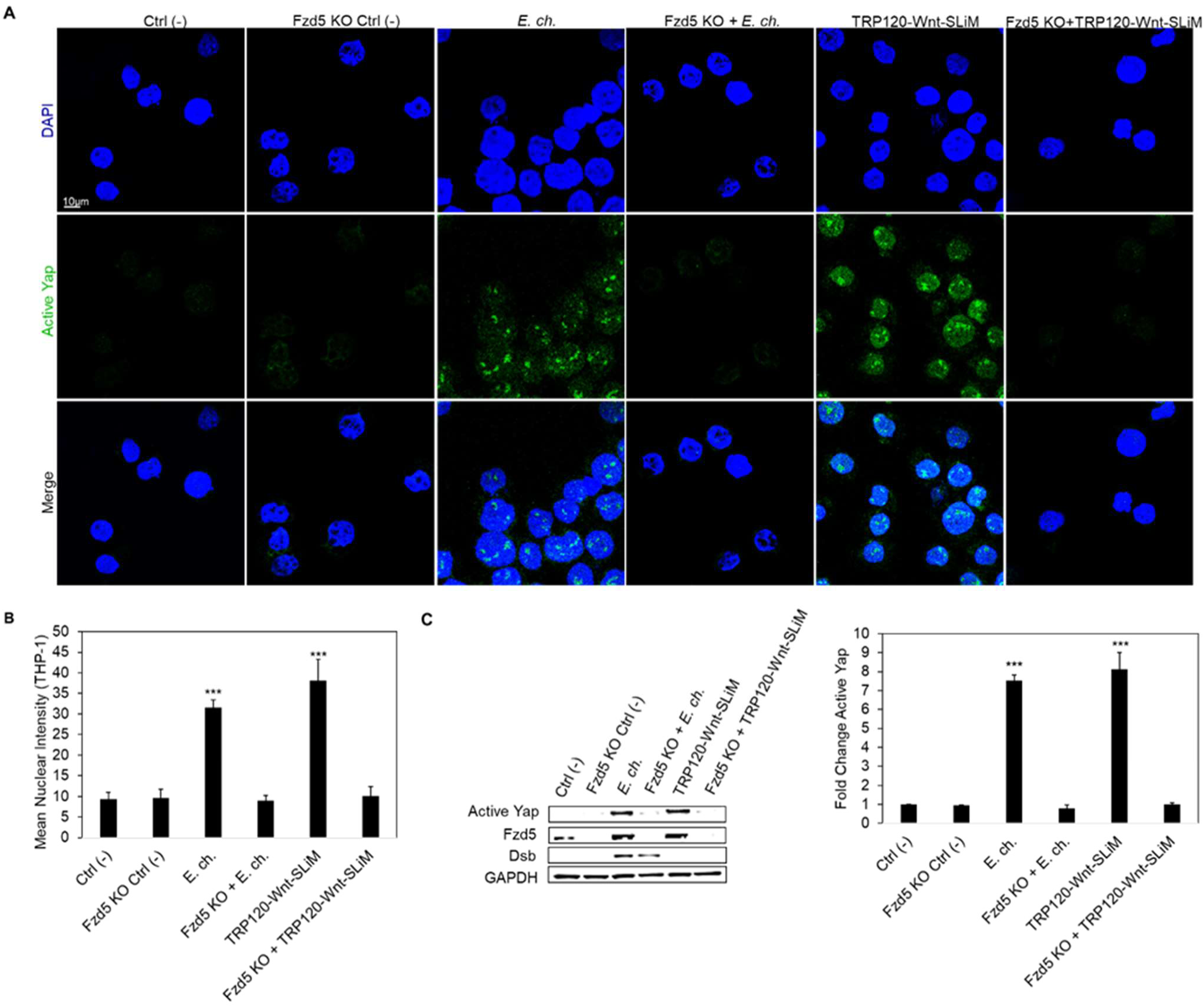
Hippo deactivation is dependent on Fzd5 receptor. (A) Confocal immunofluorescence microscopy of untreated (−), *E. chaffeensis*-infected (MOI 100) or TRP120-Wnt-SLiM treated (1 μg/mL) THP-1 cells compared to Fzd5 receptor knockout (KO) THP-1 cells. THP-1 cells and Fzd5 receptor KO THP-1 cells were harvested 6 hpt, immunostained with active Yap antibody (green), and visualized by confocal fluorescence microscopy (scale bar = 10 μm). Randomized areas/slide (n=10) were selected to detect active Yap nuclear translocation. (B) Intensity graph demonstrates the mean nuclear accumulation of active Yap in respective THP-1 cells. Analysis was performed using ImageJ to determine mean grey value from randomized areas/slide (n=10). (C) Western blot analysis of treatment groups to determine active Yap, Fzd5 and Dsb levels with GAPDH as a loading control. Western blot bar graph was analyzed from densitometry values normalized to GAPDH. (A-C) Experiments were performed with three biological and technical replicates and significance was determined through *t*-test analysis. Data are represented as means ± SD (\*\*\**p*< 0.001).

### Hippo target gene *SLC2A1* is upregulated during *E. chaffeensis* infection

To understand the basis and downstream effects of Hippo regulation during *E. chaffeensis* infection, Hippo target gene, anti-apoptotic *SLC2A1*, was investigated. *SLC2A1* encodes glucose transporter GLUT1, which is necessary in preventing apoptosis through the Yap-GLUT1-BCL-xL axis (17, 26, 29, 36, 46). During *E. chaffeensis* infection, significant upregulation of *SLC2A1* was detected at 3 and 24 hpi **(Fig. 8A)**. Further, TRP120-Wnt-SliM upregulated *SLC2A1* in a concentration dependent manner 6 hpt **(Fig 8B)**. To determine whether *E. chaffeensis* infection relies on *SLC2A1* for survival, we used RNAi to silence *SLC2A1* in THP-1 cells. Ehrlichial load was significantly reduced (24 hpi) in *SLC2A1* siRNA transfected cells compared to the scramble control transfected cells **(Fig. 8C)**. The results demonstrate that *E. chaffeensis* infection regulates and relies on *SLC2A1* expression during infection.

**Fig 8.**
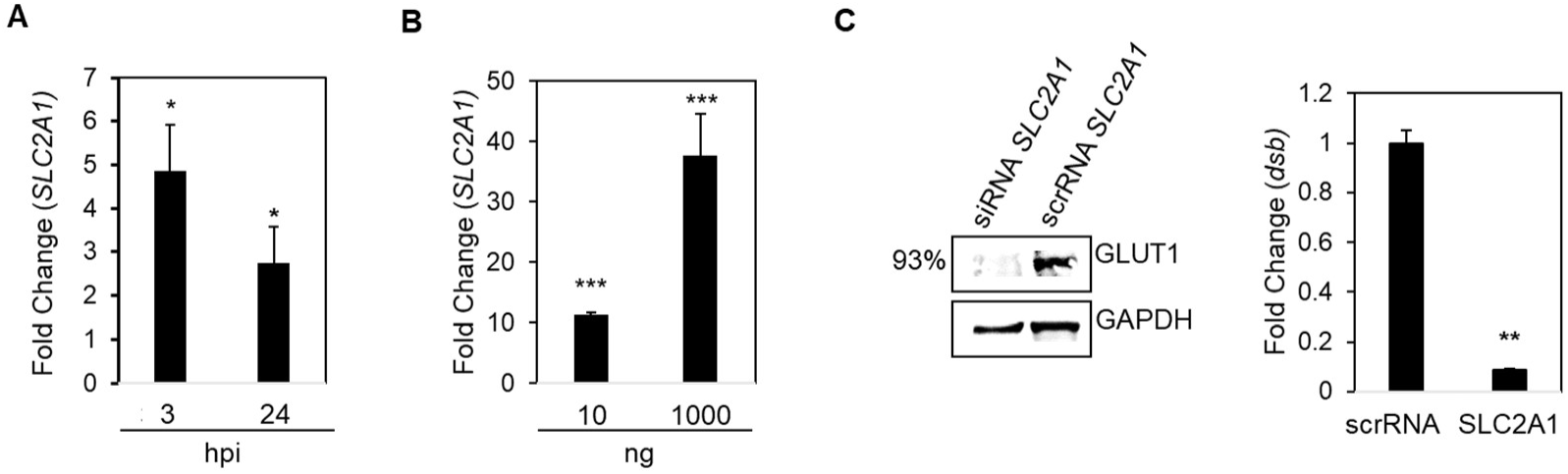
Hippo target gene *SLC2A1* is upregulated during *E. chaffeensis* infection. (A-B) Real-time qPCR analysis of anti-apoptotic regulator, *SLC2A1*, normalized to *GAPDH* during *E. chaffeensis* infection (MOI 100) at 3 and 24 hpi (A) and TRP120-Wnt-SLiM treatment (10 ng and 1000 ng) (B), demonstrating transcriptional activation. (C) Western blots depict knockdown efficiency of small interfering RNA-transfected (siRNA) THP-1 cells, with scrambled siRNA (scrRNA) transfected THP-1 cells as control from whole-cell lysates harvested at 24 h-post-transfection (as described in Fig. 5). siRNA knockdown (%) indicates total percent knockdown of *SLC2A1* relative to control, normalized to GAPDH. qPCR amplification of the ehrlichial disulfide bond formation protein (*dsb*) gene was used to quantify *E. chaffeensis* infection (MOI 100) at 24 hpi. Infected scrRNA cells are represented as positive control. (A-C) Experiments were performed with three biological and technical replicates and significance was determined through *t*-test analysis. Data are represented as means ± SD (\**p*< 0.05; \*\**p*< 0.01; \*\*\**p*< 0.001).

### *E. chaffeensis* TRP120 Wnt SliM-mediated regulation of GLUT1, BCL-xL and Bax

It is well documented that Hippo signaling promotes cell proliferation and prevents cell apoptosis through the Yap-GLUT1-BCL-xL axis (17, 26, 29, 36, 46). Further, BCL-xL is involved in the inhibition of mitochondria-mediated pro-death pathway by directly inhibiting Bax and subsequent caspase activation (46–48). Based on our results we hypothesized that *E. chaffeensis* deactivates Hippo signaling and activates Yap to increase GLUT1 and BCL-xL and decrease Bax levels. To examine this question, THP-1 cells were infected with *E. chaffeensis* or treated with TRP120-Wnt-SliM and TRP120-Wnt-SliM-mut. *E. chaffeensis* and TRP120-Wnt-SliM significantly increased GLUT1 and BCL-xL and decreased Bax levels compared to controls, consistent with a Yap-dependent anti-apoptotic profile induced by *E. chaffeensis* **(Fig. 9A-D**).

**Fig 9.**
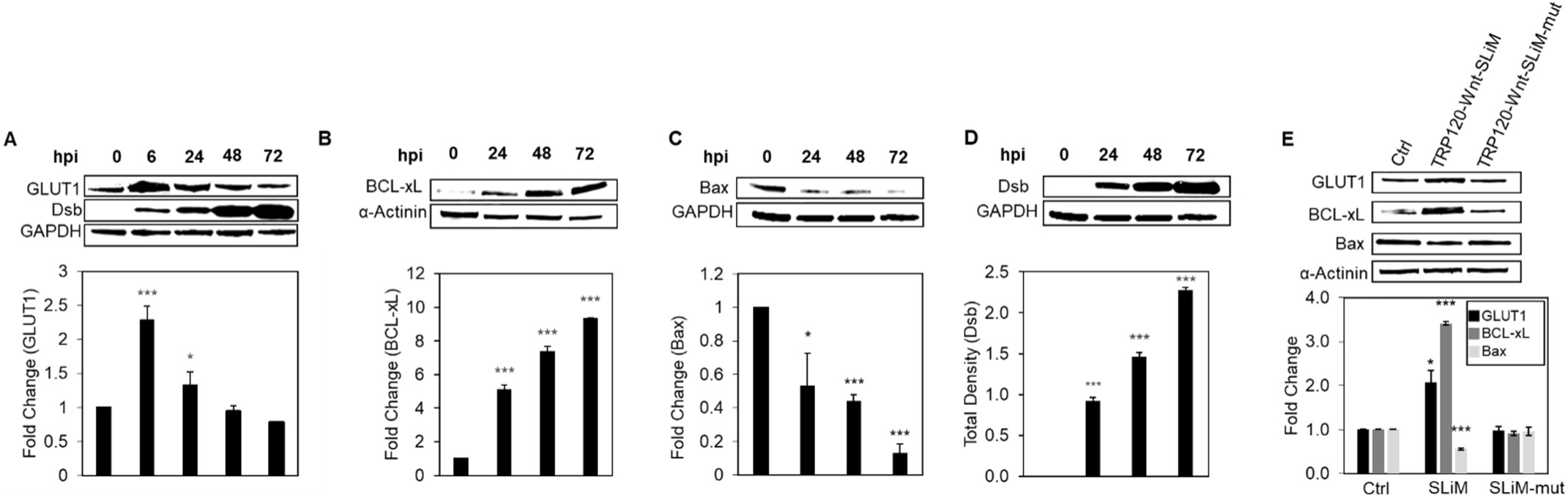
*E. chaffeensis* TRP120 Wnt SLiM-mediated regulation of GLUT1, BCL-xL and Bax. (A-C) Western blot analysis of GLUT1, BCL-xL and Bax levels during *E. chaffeensis* infection at 0, 6, 24, 48 and 72 hpi. GAPDH or α-Actinin were used as loading controls and Dsb as an infection control (D). (E) GLUT1, BCL-xL and Bax levels during TRP120-Wnt-SLiM, TRP120-Wnt-SLiM-mut-treated (1 μg/mL) and untreated THP-1 cells (24 hpt) with GAPDH as a loading control. (A-E) Bar graphs depict Western blot densitometry values normalized to GAPDH or α-actinin. Experiments were performed with three biological and technical replicates and significance was determined through *t*-test analysis. Data are represented as mean ± SD (\**p*< 0.05; \*\*\**p*< 0.001).

### TRP120 Wnt SliM-mediated regulation of GLUT1, BCL-xL and Bax during Yap inhibition

Our results support the importance of the anti-apoptotic Yap-GLUT1-BCL-xL axis during infection. Further, we hypothesized that *E. chaffeensis* deactivates Hippo to regulate GLUT1, BCL-xL and Bax. To test this hypothesis, we used a Yap inhibitor (Verteporfin) to determine whether *E. chaffeensis* infection or TRP120 Wnt SliM regulated GLUT1, BCL-xL and Bax levels during Yap inhibition. During infection, there was a significant reduction in GLUT1 in the presence of Verteporfin, demonstrating that *E. chaffeensis* depends on Yap activation to increase GLUT1. GLUT1 levels significantly increased during *E. chaffeensis* infection and in response to TRP120-Wnt-SliM treatment compared to controls **(Fig. 10B)**, consistent with results shown in **Fig. 9**. Further, during Verteporfin treatment, BCL-xL levels were unchanged during *E. chaffeensis* infection and TRP120-Wnt-SliM treated cells compared to the control **(Fig 1C)**. Conversely, Bax levels significantly increased during *E. chaffeensis* infection in the presence of Verteporfin compared to the control **(Fig. 10D)**. Notably, these results demonstrate that infection, but not SliM treatment in the presence of Verteporfin, results in significantly reduced GLUT1 levels and significantly higher Bax levels. This is likely due to induction of an apoptotic response in the monocyte in response to infection, whereas the peptide alone does not induce an apoptotic response. Based on these results, we concluded that *E. chaffeensis* TRP120 Wnt SliM-mediated regulation of GLUT1, BCL-xL and Bax levels are linked to Yap activation.

**Fig 10.**
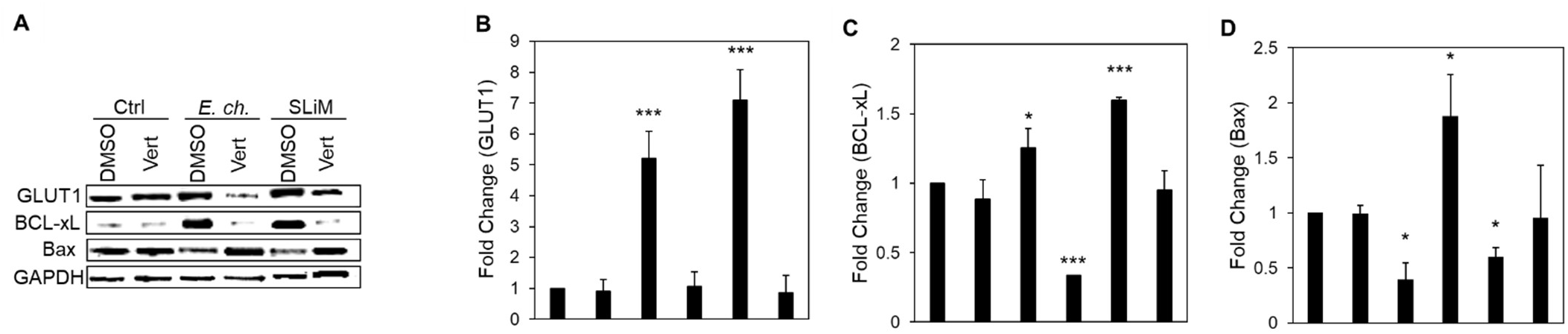
TRP120 Wnt SLiM-mediated regulation of GLUT1, BCL-xL and Bax during Yap inhibition. (A-D) Western blot analysis of GLUT1, BCL-xL and Bax levels during *E. chaffeensis* infection or TRP120-Wnt-SLiM-treatment (1 μg/mL). THP-1 cells in the presence of Yap inhibitor, Verteporfin (Vert), collected at 24 h with GAPDH as a loading control. (B) GLUT1 protein expression is significantly reduced during infection in the presence of Verteporfin compared to all groups, while GLUT1 protein expression significantly increases during *E. chaffeensis* infected and TRP120-Wnt-SLiM treated groups in the presence of DMSO (control) compared to normal cells. (C) BCL-xL level significantly increases during *E. chaffeensis-*infection and in TRP120-Wnt-SLiM treated cells in the presence of DMSO (control) compared to Verteporfin. (D) Bax levels significantly increase during infection in the presence of Verteporfin compared to DMSO (control). (B-D) Bar graphs depict Western blot densitometry values normalized to GAPDH. Experiments were performed with three biological and technical replicates and significance was determined through *t*-test analysis. Data are represented as mean ± SD (\**p*< 0.05; \*\*\**p*< 0.001).

### Yap inhibition induces an apoptotic profile during *E. chaffeensis* infection

Our results support the importance of the anti-apoptotic Yap-GLUT1-BCL-xL axis during infection and demonstrate that *E. chaffeensis* infection and TRP120-Wnt-SliM engage the Hippo pathway to regulate GLUT1, BCL-xL and Bax. Further, we hypothesized that *E. chaffeensis* regulates the anti-apoptotic Yap-GLUT1-BCL-xL axis during infection to prevent subsequent Caspase-9 and -3 activation and intrinsic apoptosis. To test this hypothesis, *E. chaffeensis*-infected and uninfected THP-1 cells were treated with Yap inhibitor Verteporfin or DMSO (**Fig. 11A**). *E. chaffeensis-*infected Verteporfin-treated cells demonstrated a significant increase in cytoplasmic condensation (precursor to apoptosis) at 24 hpi compared to uninfected Verteporfin-treated cells and *E. chaffeensis*-infected and uninfected DMSO-treated cells, supporting the conclusion that *E. chaffeensis* activates Yap to prevent apoptosis. Additionally, ehrlichial survival was significantly reduced in the presence of Verteporfin compared to the control (DMSO) **(Fig. 11B)**. Further, cell viability significantly decreased in *E. chaffeensis-*infected cells treated with Verteporfin **(Fig. 11C)**. To define a direct mechanism by which *E. chaffeensis* activates Yap to prevent apoptosis, we evaluated levels of pro and cleaved Caspase-9 and -3 during infection in the presence of Verteporfin **(Fig. 11D-E)**. *E. chaffeensis-*infected cells treated with inhibitor showed a significant decrease in pro-Caspase-9 and -3 levels, while cleaved Caspase-9 and -3 levels significantly increased during *E. chaffeensis* infection in the presence of Verteporfin compared to DMSO-treated *E. chaffeensis*-infected cells (control). Collectively, these results define a mechanism whereby *E. chaffeensis* activation of Yap regulates Caspase-9 and -3 to inhibit intrinsic apoptosis.

**Fig 11.**
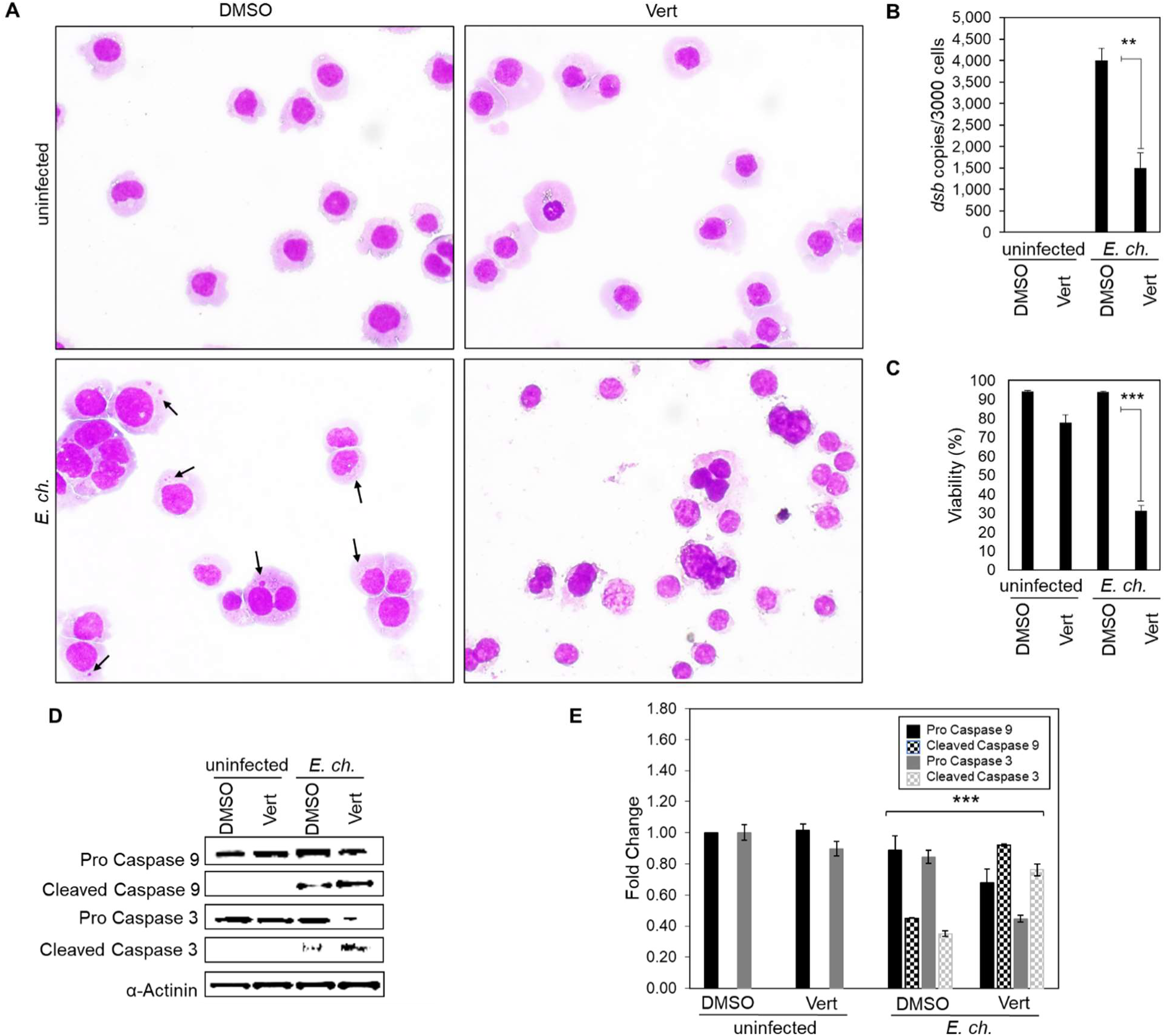
Yap inhibition induces an apoptotic profile during *E. chaffeensis* infection. (A) Brightfield micrographs showing effects of DMSO or Hippo inhibitor Verteporfin on uninfected and *E. chaffeensis*-infected THP-1 (MOI 50) cells prepared using Diff-Quick staining. *E. chaffeensis*-infected or uninfected THP-1 cells were treated with DMSO or Verteporfin (7 μg/mL) and collected 24 h later. *E. chaffeensis-*infected THP-1 cells treated with Verteporfin undergo cytoplasmic condensation (precursor to apoptosis), but other treatment groups do not (arrows point to morulae). (B) Bar graph showing fold-change in *E. chaffeensis* infection for each treatment group. Ehrlichial loads were determined using qPCR measurement of *dsb* copy and normalized to *GAPDH. E. chaffeensis* infection significantly declines in the presence of Verteporfin. (C) Bar graphs showing cell viability for each treatment group. Cell viability was determined using the Cellometer Mini bright field automated cell counter and the pattern-recognition assay. Cell viability significantly declines in the presence of Verteporfin during *E. chaffeensis* infection. (D-E) Western blot analysis of Caspase-9 and -3 levels for each group with α-actinin as a loading control. Pro-Caspase-9 and -3 levels significantly decreased while cleaved Caspase-9 and -3 levels significantly increased during *E. chaffeensis* infection in the presence of Verteporfin compared to DMSO control. Bar graphs depict Western blot densitometry values normalized to α-actinin. Experiments were performed with three biological and technical replicates and significance was determined through *t*-test analysis. Data are represented as means ± SD (\**p*<0.05; \*\**p*<0.01; \*\*\**p*<0.001).

## Discussion

In 2003, the Hippo pathway was discovered and has since been recognized as a key pathway that regulates embryogenesis, organ size, tissue homeostasis, cell proliferation, apoptosis, and tumorigenesis (14–18). Recently, investigations have linked pathway crosstalk between the Wnt and Hippo signaling pathways to control cell fate, demonstrating that Wnt5a and Wnt3a ligands bind the Fzd5 receptor to deactivate Hippo signaling and activate Yap (10, 18, 21). Additional crosstalk occurs between Yap and Wnt transcriptional regulator, β-catenin. When Hippo is active, Yap is phosphorylated and remains within the cytoplasm where it sequesters β-catenin, leading to the degradation of β-catenin and inhibition of Wnt signaling (18). Thus, it is imperative that Hippo deactivation by interactions between Wnt5a/Wnt3a and Fzd5 receptor occur to support Wnt signaling. Notably, *E. chaffeensis* is a known β-catenin activator and utilizes a TRP120 Wnt SliM to activate β-catenin for Wnt gene regulation (6). Based on the premise of Hippo-Wnt crosstalk and the regulation of Wnt signaling by the TRP120 Wnt SLiM, we sought to identify whether the TRP120 Wnt SLiM deactivates Hippo leading to Yap activation. Indeed, we reveal that the TRP120 Wnt SliM regulates Hippo signaling and identified the downstream effects directed at inhibiting intrinsic host-cell apoptosis. *Ehrlichia chaffeensis* contains a Wnt SliM and depends on the Wnt Fzd5 receptor to activate transcription co-activator Yap, which promotes a significant upregulation in genes critical for Hippo and Wnt signaling. This is the first report of a single eukaryotic SliM mimetic in bacteria that can regulate multiple conserved signaling pathways, which reveals a novel strategy utilized by obligate intracellular bacteria to extend host cell lifespan and highlights the importance of pathogen utilization of eukaryotic cellular signaling motifs for reprogramming the host cell to promote infection.

Although Hippo signaling has been studied during viral infection, little is known regarding Hippo signaling during bacterial infection. We investigated whether *E. chaffeensis* regulates Hippo signaling during infection. Indeed, we confirmed that infection induces Yap activation and transcriptional induction of Hippo pathway genes including crucial components of the Hippo and Wnt signaling pathways, *YAP*, *TAZ*, *TEAD1*, *TEAD2, TEAD3, TEAD4,* and *DVL2.* Further, we determined that TRP120 induces Yap activation and subsequent Hippo gene regulation, including Hippo and Wnt targets *YAP, TAZ, TEAD4,* and *DVL2*, similarly to Wnt5a, which significantly upregulated *YAP*, *TAZ*, *TEAD1*, *TEAD2, TEAD3, TEAD4, Wnt1* and *DVL2*. Although there was differential expression of Hippo pathway genes in TRP120 and Wnt5a-treated cells, we discovered that many Hippo target genes were upregulated by both TRP120 and Wnt5a, which supports TRP120 mimicry of Wnt5a. Some differences between TRP120 and Wnt5a were expected since biological functions between various Wnt ligands differ despite a highly similar amino acid sequence (49). Additionally, TRP120 also contains Notch and Hedgehog SliMs which may also influence gene expression due to the intricate crosstalk between the pathways (50, 51).

To further establish the direct mechanism of Hippo regulation during infection, we determined that the TRP120-Wnt-SliM sufficiently induces active Yap. Notably, *E. chaffeensis,* TRP120 and TRP120-Wnt-SliM induced similar Yap activity in THP-1 cells and primary human monocytes, which is important to note since primary cells have a limited lifespan and THP-1 cells are a more practical alternative for laboratory studies. Additionally, we further defined the previously reported Wnt SliM (6), shortening the SliM to 6 aa (from 17 aa) using BLAST analysis to detect a short region of homology among TRP120 and Wnt5a/3a ligands. The shorter TRP120-Wnt-SliM highlights shared amino acids between Wnt5a/3a that may be critical in binding Fzd receptors and activating signaling. In our previous study, we defined the TRP120-Wnt-SliM based on sequence and functional similarities between TRP120 and Wnt8, since Wnt8 activates β-catenin and structural studies have defined many residues for Wnt8-Fzd binding (6, 52). However, many of the Wnt residues necessary for binding Fzd receptors are not conserved amongst the Wnt ligands (53). Additionally, Wnt5a and Wnt3a residues for Fzd binding are not well defined; however, these ligands are relevant to this study since they activate Yap (10). Identification of a SliM with capability to affect multiple pathways is new to science and will have significant impact in how these ligand receptor interactions are viewed by cell biologists and others.

In our investigation, TRP120-Wnt-SliM exhibited stronger upregulation of Hippo gene targets than TRP120. This is likely due to higher molar concentrations of SliM sequence present in the TRP120-Wnt-SliM treatment. Nevertheless, Hippo gene regulation profiles were similar between *E. chaffeensis,* TRP120, Wnt5a and TRP120-Wnt-SliM. To further support our results, we used the Wnt SliM (QDVASH) to target Wnt signaling and determined that it does activate both Hippo and Wnt signaling, consistent with known Hippo-Wnt receptor overlap and crosstalk (18). Additionally, we used an anti-SliM antibody which blocked Yap activation during *E. chaffeensis* infection and TRP120-Wnt-SliM treatment, demonstrating the importance of the SliM in Hippo regulation during infection and confirmed that the TRP120-Wnt-SliM is the only SliM mimetic utilized by *E. chaffeensis* to activate Yap.

In recent years, our laboratory has determined that TRP120 contains multiple SliMs within the intrinsically disordered TRD that act as ligand mimetics to regulate Wnt, Notch, Hedgehog and Hippo signaling. *E. chaffeensis* likely contains multiple pathway activating SliMs due to the intricate crosstalk between the pathways, and the role each plays in regulating apoptosis to promote infection (18, 51, 54). SliMs are disordered, short, linear sequence and contain a limited number of specificity-determining residues (55). Few mutations are necessary for the generation of new SliMs, allowing rapid convergent evolution of SliMs within proteins *de novo*, enabling rapid functional flexibility (56, 57). *E. chaffeensis* has likely convergently evolved TRP120 SliMs to engage multiple cellular signaling pathways for redundancy and to influence anti-apoptotic signaling through different pathways. All defined TRP120 SliMs activate conserved signaling pathways known to prevent apoptosis, which may be a strategy executed by *E. chaffeensis* to insure host cell survival and productive infection.

TRP120 is a Wnt ligand mimic and directly interacts with Fzd5 receptor (6). Wnt5a and Wnt3a ligands interact with Fzd5 receptor, which can lead to activation of Hippo and Wnt transcriptional regulators Yap and β-catenin, respectively. Further, while only Fzd −1, −2 and −5 are associated with Yap activation, most Fzd receptors are known to activate β-catenin (6, 10, 24, 44). Additionally, the co-expression of Fzd5 with co-receptor tyrosine kinase ROR1 potentiates Fzd5 receptor-induced Yap activation (10). Previously, we demonstrated that *E. chaffeensis* survival depends on ROR1, which may be due to its role in co-activation of Yap (6, 10). To better understand why *E. chaffeensis* interacts with the Fzd5 receptor and how it relates to Yap activation, we utilized Fzd5 receptor KO to demonstrate that Fzd5 receptor is essential for Yap activation during infection. We found that *E. chaffeensis* and TRP120-Wnt-SliM Yap activation is solely dependent on the Fzd5 receptor. Yap activation has been associated with Fzd −1, −2 and −5 in HEK293 cells (10, 21); however, the fact that Yap activation induced by *E. chaffeensis* and TRP120-Wnt-SliM depends solely on the Fzd5 receptor may be related to fundamental differences in cell types (innate immune phagocyte vs. epithelial kidney cell). In contrast to our finding that Hippo relies completely on the Fzd5 receptor for signaling, β-catenin activation was only significantly reduced (∼50%) in the Fzd5 receptor knockout cells. This is consistent with reports demonstrating the multiple Wnt ligands and Fzd receptors are involved in β-catenin activation. Similarly, we have observed interactions between TRP120 and other Fzd receptors known to activate β-catenin (6).

Cellular apoptosis plays an important role as an innate defense mechanism against microbial infection. During infection, cells utilize apoptotic mechanisms for processing infected apoptotic bodies containing pathogens to facilitate antigen presentation and protective immunity (58). Preventing apoptosis is critical to obligately intracellular bacteria since maintaining a replicative niche is essential to complete the infection cycle. Obligate intracellular pathogens including *Rickettsia, Anaplasma, Mycobacterium, Chlamydia,* and others have evolved multiple regulatory mechanisms to inhibit host cell apoptosis, including regulation of mitochondria-mediated intrinsic apoptosis (58–65). Additionally, intracellular bacteria regulate the BCL-2 family of proteins to stabilize mitochondria to promote host cell survival. Recently, we demonstrated that *E. chaffeensis* activates the Hedgehog pathway to regulate mitochondria-mediated intrinsic apoptosis via BCL-2 and extend the host cell lifespan (9). *Chlamydia trachomatis* upregulates MCL-1 to inhibit Bax-induced apoptosis (66), and *M. tuberculosis* upregulates BCL-2 in macrophages during infection to prevent apoptosis (67).

Recently, investigations have demonstrated a major role for Hippo signaling in glucose metabolism to preserve mitochondria stabilization and prevent apoptosis. To prevent apoptosis, the cell deactivates Hippo signaling to activate the transcriptional co-activator Yap to upregulate Hippo gene targets including *SLC2A1*, which encodes glucose transporter GLUT1. The upregulation of GLUT1 promotes glucose metabolism, which subsequently promotes the upregulation of BCL-xL (26, 29–31). Previous studies demonstrate that a reduction in GLUT1 protein expression increases Bax, Bak, Bim and Bid (pro-apoptotic) and inhibits MCL-1 and BCL-xL (36). Additionally, *E. chaffeensis* infection and TRP120-Wnt-SliM treatment increased GLUT1 and BCL-xL and decreased Bax levels. Further, we show that a small molecule Yap inhibitor prevents *E. chaffeensis* from regulating GLUT1, BCL-xL, and Bax, and induces a pro-apoptotic profile. These results reveal a novel anti-apoptotic mechanism by which *E. chaffeensis* modulates the Hippo pathway for infection by extending the host cell lifespan using glucose metabolism, which is consistent with the role of Hippo signaling in cell biology. Remarkably, *Ehrlichia chaffeensis* regulates Hippo and Hedgehog to target various BCL-2 family proteins and inhibit intrinsic apoptosis, a remarkable redundancy resulting in comprehensive regulation of anti-apoptotic signaling for intracellular survival.

The current study reveals a model of eukaryotic mimicry where a single bacterial SLiM phenocopies endogenous ligands to regulate multiple conserved signaling pathways. Here, we characterize a TRP120 Wnt SLiM that utilizes Hippo-Wnt pathway crosstalk to engage the Yap-GLUT1-BCL-xL axis to promote an anti-apoptotic profile (**Fig. 12)**. This study demonstrates the importance of Hippo signaling in preventing apoptosis for ehrlichial replication and provides a potential new target for therapeutic development. The potential to use *E. chaffeensis* as a model to define the role of SLiM ligand mimicry and evolutionary conserved eukaryotic signaling pathway will lead to a broader understanding of intracellular pathogen biology and provide mechanistic targets for countermeasure development.

**Fig. 12.**
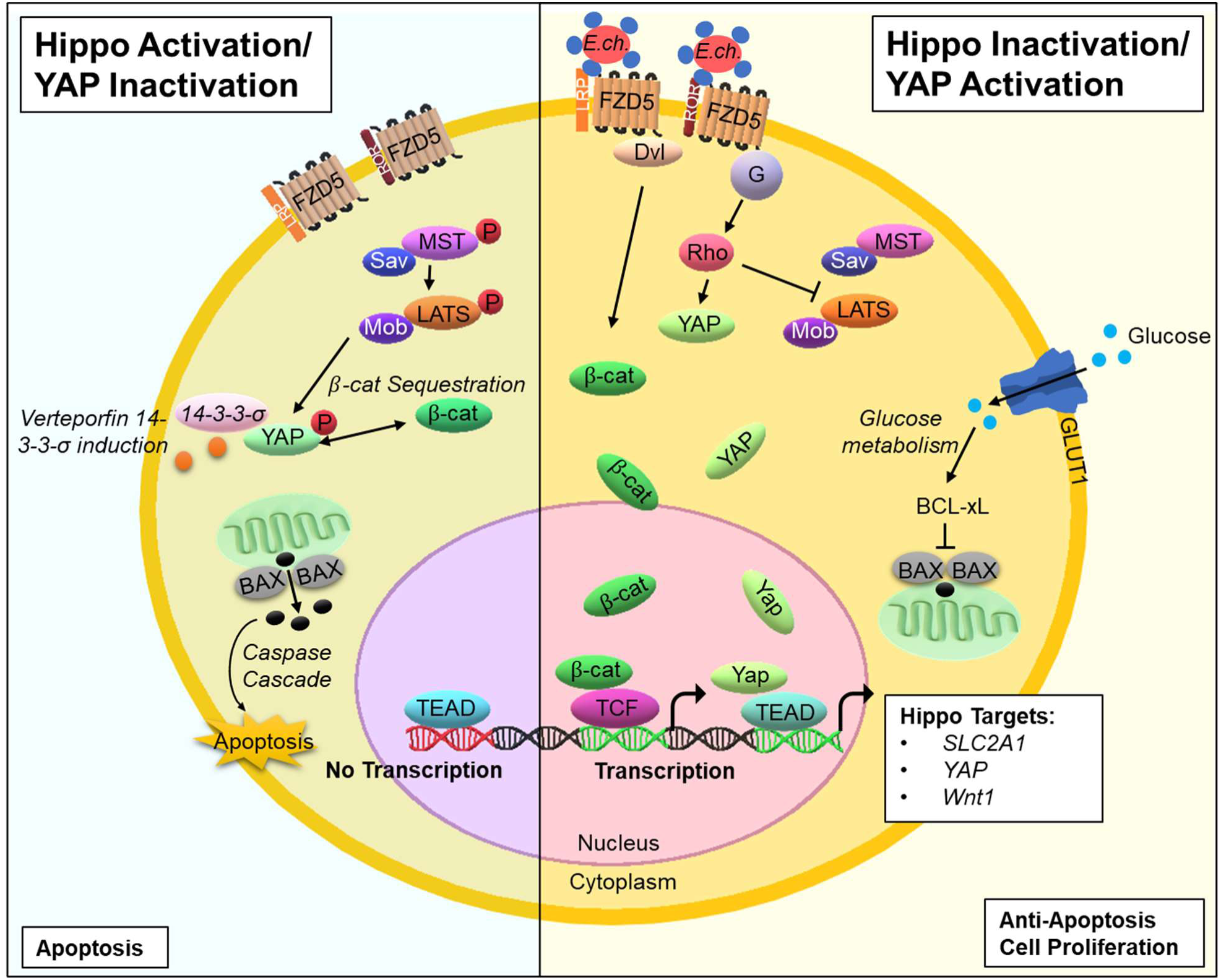
Model of *E. chaffeensis* TRP120 mediation of Hippo signaling and downstream maintenance of signaling. Activation of Hippo signaling mediates cell fate through the phosphorylation and deactivation of Yap, which further leads to β-catenin deactivation and host cell apoptosis. In response, Wnt ligands bind the Fzd5 receptor at the cysteine-rich extracellular domain (ECD) to activate Yap nuclear translocation and promote β-catenin nuclear translocation, making Yap a potential target of the TRP120 Wnt SLiM. Therefore, DC ehrlichia surface-expressed TRP120 directly engages the Fzd5 receptor at the extracellular conserved cysteine-rich domain through a Wnt SLiM repeated in the TRP120 TRD, thus activating transcription co-activators Yap and β-catenin to translocate freely to the nucleus to bind DNA and upregulate Hippo gene targets. Further, Yap nuclear translocation leads to the upregulation of target gene *SLC2A1*, which encodes GLUT1 and therefore induces glucose metabolism to prevent host cell apoptosis via BCL-xL inhibition of Bax.

## Materials and Methods

### Cell culture and *E. chaffeensis* cultivation

Human monocytic leukemia cells (THP-1; ATCC TIB-202) or primary human monocytes (PHMs) were propagated in RPMI 1640 with L-glutamine and 25 mM HEPES buffer (Invitrogen, Carlsbad, CA), supplemented with 10% fetal bovine serum, and incubated at 37°C in a 5% CO_2_ atmosphere. Peripheral blood mononuclear cells were obtained from deidentified healthy human donors (Gulf Coast Regional Blood Center, Houston, TX) and primary human monocytes isolated using MACS negative selection (Miltenyi Biotec, Cambridge, MA). *E. chaffeensis* (Arkansas strain) was cultivated in THP-1 cells and primary human monocytes as previously described (9).

### Protein sequence analysis

The NCBI Protein Basic Local Alignment Search Tool (Protein BLAST) was utilized for sequence alignment of TRP120 (NCBI accession number AAO12927.1) and Wnt5a and Wnt3a amino acid sequences (NCBI accession numbers AAH74783 and EAW69829).

### Recombinant proteins and peptides

*E. chaffeensis* recombinant full length TRP120 (rTRP120-FL), TRP120 TRD (rTRP120-TR) or thioredoxin (rTrx; ctrl) were expressed in *E. coli* and purified as described previously (8). rWnt5a (R&D Systems, Minneapolis, MN) and peptides (GenScript, Piscataway, NJ) were obtained from a commercial source. Synthesized peptides include TRP120-TR-Wnt5a (IKDLQDVASHESGVSDQPA; represents the entire homologous Wnt5a sequence), TRP120-TR (−) (SHQGETEKESGITESHQKEDEI; neg ctrl), TRP120-Wnt-SLiM (QDVASH), TRP120-Wnt-SLiM-mut (IKDLGAGAGAESGVS; Gly/Ala substitutions in the Wnt SLiM motif) and TRP120-Wnt-QDVAS (QDVAS).

### Antibodies and inhibitors

Antibodies used in this study include α-disulfide bond formation protein (Dsb) (68), α-TRP120-Wnt-SLiM (targets TRP120 sequence DL<u>QDVASH</u>ESGVSDQPAQV)(6), α-TRP120 (6), α-active Yap (Abcam, Cambridge, UK), α-Yap, α-TEAD(1, 3, and 4)(Santa Cruz Biotechnology, Dallas, TX), α-active β-catenin, α-Fzd5 receptor, α-BCL-xL, α-Bax, α-Caspase-3, and -9 (Cell Signaling, Danvers MA), α-GLUT1 (Abcam, Cambridge, UK), α-GAPDH (MilliporeSigma, Burlington, MA), α-rabbit IgG (H+L) Alexa Fluor Plus 594 and α-mouse IgG (H+L) Alexa Fluor Plus 488 (Invitrogen Carlsbad, CA). Inhibition of the Hippo pathway was performed using Verteporfin (Thermo Fisher Scientific, Waltham, MA).

### Neutralization assay

*E. chaffeensis* or TRP120-Wnt-SLiM were incubated for 1 h or overnight, respectively, with 1.5 μg/mL of either α-TRP120-Wnt-SLiM antibody (targets TRP120 sequence SKVEQEET<u>NPEVLIKD</u>LQDVAS) or α-TRP120-PIS antibody (control), and then THP-1 cells were subsequently treated with each mixture for 10 h.

### RNAi and *Ehrlichia* quantification

All siRNAs were ON-TARGETplus SMARTpool (Dharmacon, Lafayette, Co). siRNA KD was performed as previously described (6, 9). Scrambled RNAi was used as siRNA control. THP-1 cells were infected with cell-free *E. chaffeensis* (MOI 100) 24 h post-transfection. Cells were harvested at 24 hpi and ehrlichial load was determined using qPCR as previously described (72). All knockdowns were performed with three biological and technical replicates and significance was determined using a *t*-test analysis.

### Confocal microscopy

*E. chaffeensis-*infected (MOI 100) and uninfected THP-1 cells were seeded in T-150 flasks (Corning, Lowell, MA) at 30% confluency and collected at 0, 4, 10, 24 and 48 hpi. THP-1 cells were treated with rTRP120-FL, rTrx (−), rWnt5a, TRP120-TR-Wnt5a, TRP120-TR (−), TRP120-Wnt-SLiM or TRP120-Wnt-SLiM-mut (1 μg/mL) and collected 6 hpt for confocal microscopy as previously described (9). *E. chaffeensis-*infected, uninfected, rTRP120-FL, rTrx (−), rWnt5a, and TRP120-Wnt-SLiM and TRP120-Wnt-SLiM-mut peptide-treated primary human monocytes were seeded at 30% confluency in 12-well plates (Corning) containing a coverslip and incubated for 10 h. Cells were prepared for confocal microscopy as previously described (9) and stained with mouse anti-active Yap monoclonal antibody (1:200), rabbit anti-active β-catenin monoclonal antibody (1:100) and rabbit anti-Dsb antibody (1:500). Secondary antibodies were α-rabbit IgG (H+L) Alexa Fluor Plus 594 and α-mouse or rabbit IgG (H+L) Alexa Fluor Plus 488 (1:200). Zeiss LSM 880 laser microscope was utilized to obtain all confocal laser micrographs and analyzed with Zen black and Fiji software. Randomized areas/slide (n=10) were used to detect active Yap. Experiments were performed with three biological and technical replicates.

### RNA isolation and cDNA synthesis

*E. chaffeensis*-infected (MOI 100), uninfected, rTRP120-FL, rTrx (−), rWnt5a, TRP120-Wnt-SLiM (10 ng/mL or 1 μg/mL) cells were harvested at 24 h. Uninfected/untreated or rTrx (−)-treated cells were used as controls for infection and protein/peptide treatments. RNA isolation and cDNA synthesis was performed as previously described (9). Data was generated from three biological and technical replicates.

### Human Hippo signaling pathway PCR array

The human Hippo signaling target PCR array (Qiagen) was used to determine expression of 84 key Hippo target genes. PCR arrays were performed according to the manufacturer’s protocol (Qiagen). Real-time PCR was performed using RT^2^ Profiler PCR array, SYBR green master mix (Qiagen) using the QuantStudio 6 Flex real-time PCR system (Thermo Fisher Scientific). PCR data analysis was performed as previously described (6).

### Western immunoblot

Briefly, THP-1 cells (100% confluent) were harvested and lysates prepared using CytoBuster protein extraction reagent (Novagen/EMD, Gibbstown, NJ) supplemented with complete mini EDTA-free protease inhibitor (Roche, Basel, Switzerland) and phenylmethene-sulfonylfluoride PMSF (10 mM) (Sigma-Aldrich). Cell lysate protein concentrations were determined and Western blots were performed as previously described (9) using α-Yap, α-TEAD(1, 3 and 4), α-Fzd5 receptor, α-BCL-xL, α-Bax antibodies (1:200), α-Caspase 3 and -9 antibodies (1:100) and α-GAPDH (1:10,000). Experiments were performed with three biological and technical replicates and significance determined by *t*-test analysis.

### Fzd5 receptor knockout cells

CRISPR/Cas9 Fzd5 receptor KO THP-1 cells were obtained from a commercial source (Synthego, Redwood City, California) and serially diluted to isolate a clonal population. Normal and Fzd5 receptor KO THP-1 cells (30% confluent) were infected with *E. chaffeensis* (MOI 100) or treated with rWnt5a, TRP120-Wnt-SLiM or TRP120-Wnt-SLiM-mut (1µg/mL) and collected at 6 h for confocal and Western blot analysis.

### Real-time qPCR

The analysis of *SLC2A1* gene expression during infection was determined using RT-qPCR. THP-1 cells (100% confluent) were infected with *E. chaffeensis* (MOI 100). Cells were harvested at 0, 3 and 24 hpi to examine gene expression during the entry and early replication phase. The fold change in *SLC2A1* from 0 to 3 or 24 hpi was calculated using the 2^−ΔΔ^*^CT^* method and *C_T_* values for host *SLC2A1* and *GAPDH* genes as previously described (69).

### Hippo inhibitor infection analysis

*E. chaffeensis*-infected (MOI 50), uninfected, TRP120-Wnt-SLiM- and TRP120-Wnt-SLiM-mut-treated THP-1 cells (30% confluent) were incubated with DMSO or Verteporfin (7 μg/mL) for 24 h, then cells were harvested for Western blot and Diff-Quik staining (Thermo Fisher Scientific). Ehrlichial load was determined using qPCR as described above. Cell counts and viability were determined by the Cellometer mini brightfield automated cell counter (Nexcelom, Lawrence, MA).

## Acknowledgments

We thank Maxim Ivannikov and the Optical Microscopy Core Laboratory for assistance with confocal microscopy. This work was supported by the National Institute of Allergy and Infectious Disease grants AI137779 and AI149136 to J.W.M., T32AI007526-20 Biodefense Training Program predoctoral fellowship and Kempner Postdoctoral Fellowship Award to C.D.B., McLaughlin Endowment and NIH 1F31AI152424 predoctoral fellowships to L.L.P, Sealy Center for Vector Borne and Zoonotic Diseases and McLaughlin predoctoral fellowships to N.A.P, and T32AI007526-22 Biodefense Training Program predoctoral fellowship to R.N.S.

---

**Fig. S1.**
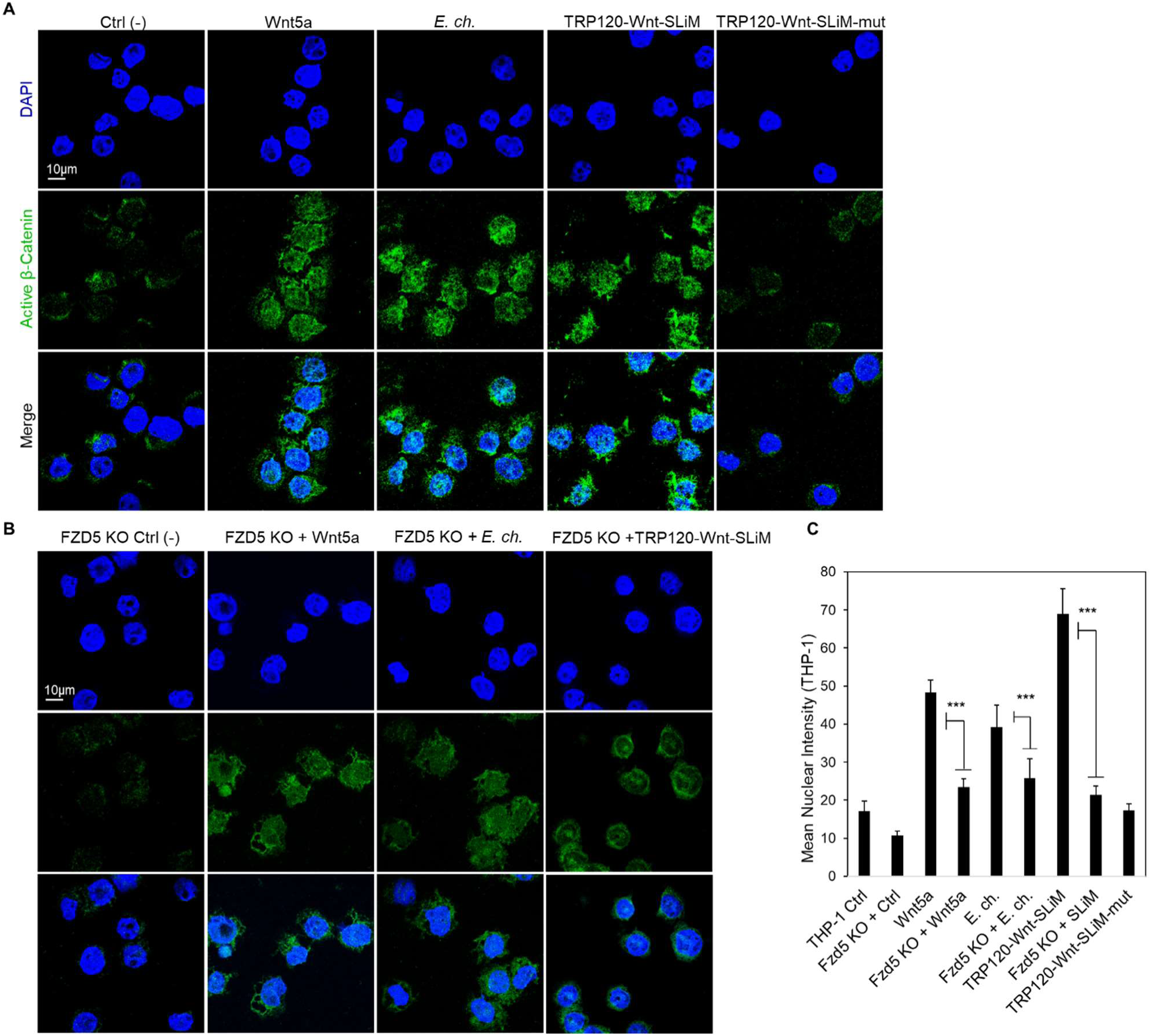
TRP120 Wnt SLiM activates Wnt signaling via FZD5 receptor. (A-B) Confocal immunofluorescence microscopy of *E. chaffeensis*-infected (MOI 100) or SLiM treated (1 μg/mL) THP-1 cells compared to untreated (−) and Wnt5a-treated (+) THP-1 cells stained with active β-catenin antibody. (A) The micrograph shows increased levels of active β-catenin (green) in Wnt5a (+), infected, and TRP120-Wnt-SLiM-treated, but not in TRP120-Wnt-SLiM-mut-treated THP-1 cells (6 hpt)(scale bar = 10 μm). (B) Confocal immunofluorescence microscopy of the Fzd5 receptor knockout (KO) THP-1 Fzd5 receptor KO cells were harvested (6 hpt) and immunostained with active β-catenin antibody (green). (A-B) Experiments were performed with three biological and technical replicates. Randomized areas/slide (n=10) were used to detect active β-catenin nuclear translocation. (C) Intensity graphs demonstrate the mean nuclear accumulation of active β-catenin in respective THP-1 cells. THP-1 FZD5 receptor KO cells have significantly less β-catenin. Analysis was performed using ImageJ and determining mean grey value from randomized areas/slide (n=10). Data are represented as means ± SD (\*\*\**p*< 0.001).

**Fig. S2.**
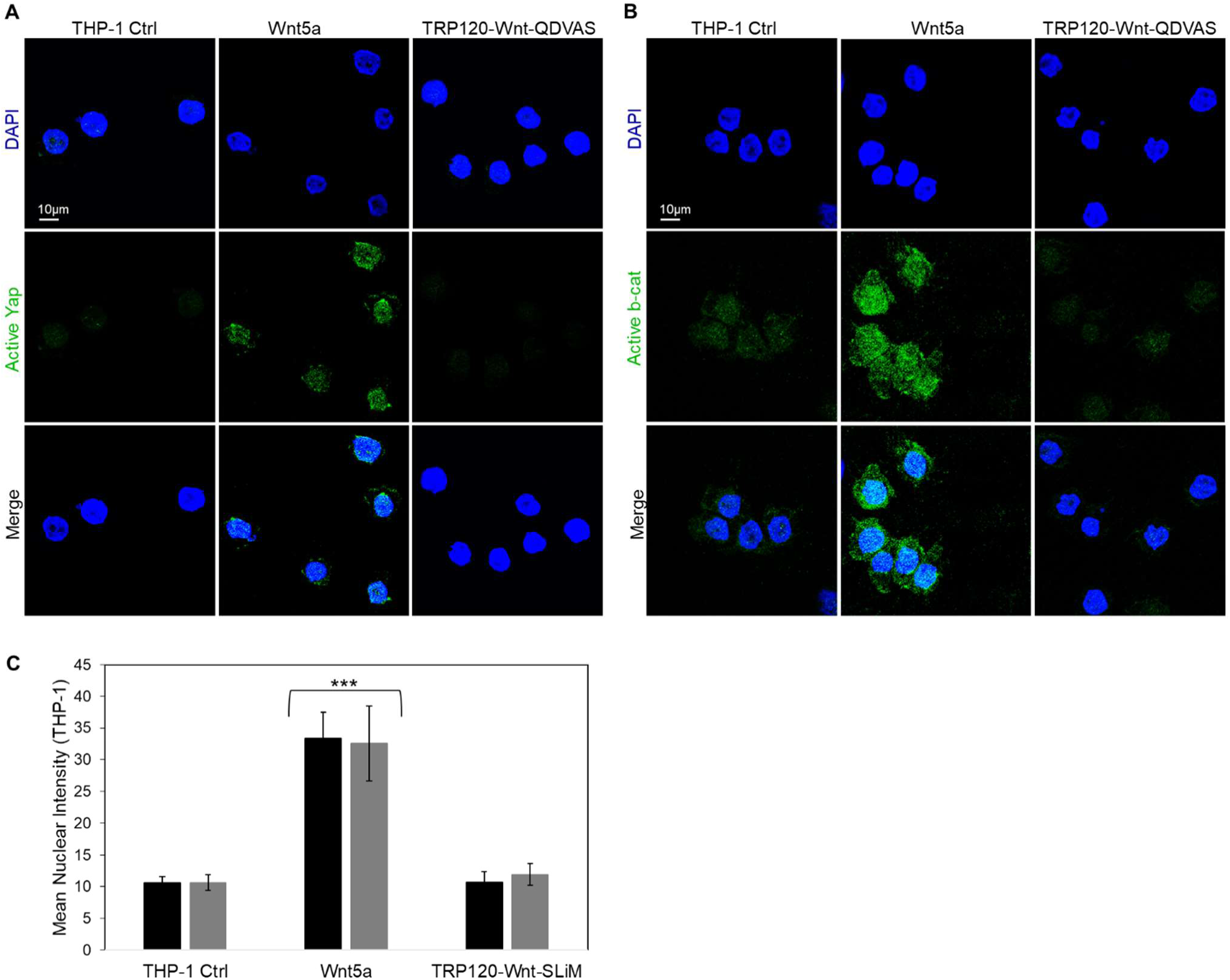
TRP120 Wnt SLiM deletion mutant does not regulate Yap or β-catenin. (A-B) Confocal immunofluorescence microscopy of TRP120-Wnt-SLiM His deletion mutant (TRP120-Wnt-QDVAS) peptide-treated (1 μg/mL) THP-1 cells compared to untreated (−) and Wnt5a-treated (+) THP-1 cells and stained with active Yap or β-catenin antibody. The micrograph shows no significant change in active Yap or β-catenin levels in TRP120-Wnt-SLiM His deletion mutant-treated cells compared to untreated (−) THP-1 cells (6 hpt)(scale bar = 10 μm). Experiments were performed with three biological and technical replicates. Randomized areas/slide (n=10) were used to detect active Yap or β-catenin nuclear translocation. (C) Intensity graphs demonstrate the mean nuclear accumulation of active Yap or β-catenin in respective THP-1 cells. Analysis was performed using ImageJ and determining mean grey value from randomized areas/slide (n=10). Data are represented as means ± SD (\*\*\**p*< 0.001).

